# PhyloAcc-GT: A Bayesian method for inferring patterns of substitution rate shifts and associations with binary traits under gene tree discordance

**DOI:** 10.1101/2022.12.23.521765

**Authors:** Han Yan, Zhirui Hu, Gregg Thomas, Scott V. Edwards, Timothy B. Sackton, Jun S. Liu

**Affiliations:** Department of Statistics, Harvard University; Informatics Group, Harvard University; Department of Organismic and Evolutionary Biology, Harvard University

**Author notes:** These authors contributed equally (Han Yan, Zhirui Hu, Gregg Thomas).

## Abstract

An important goal of evolutionary genomics is to identify genomic regions whose substitution rates differ among lineages. For example, genomic regions experiencing accelerated molecular evolution in some lineages may provide insight into links between genotype to phenotype. Several comparative genomics methods have been developed to identify genomic accelerations between species, including a Bayesian method called PhyloAcc, which models shifts in substitution rate in multiple target lineages on a phylogeny. However, few methods consider the possibility of discordance between the trees of individual loci and the species tree due to incomplete lineage sorting, which might cause false positives. Here we present PhyloAcc-GT, which extends PhyloAcc by modeling gene tree heterogeneity to detect rate shifts across genomic regions. Given a species tree, we adopt the multispecies coalescent model as the prior distribution of gene trees, use Markov chain Monte Carlo (MCMC) for inference, and design novel MCMC moves to sample gene trees efficiently. Through extensive simulations, we show that PhyloAcc-GT outperforms PhyloAcc and other methods in identifying target-lineage-specific accelerations and detecting complex patterns of rate shifts, and is robust to specification of population size parameters. We apply PhyloAcc-GT to two examples of convergent evolution: flightlessness in ratites and marine mammal adaptations. PhyloAcc-GT is usually more conservative than PhyloAcc in calling convergent rate shifts because it identifies more accelerations on ancestral than on terminal branches. In summary, PhyloAcc-GT is a useful tool to identify shifts in substitution rate associated with specific target lineages while accounting for incomplete lineage sorting.

## Introduction

The ongoing deluge of whole-genome sequences across the tree of life, combined with new phylogenetic methods, have provided comparative biologists with powerful opportunities for a detailed understanding of variable rates of change among genes and lineages, with the aim of identifying regions of the genome evolving by natural selection and potentially linked to phenotypic evolution. Differences between the sequences and structure of genomes allow us to quantify rates of change for various types of mutations and to formulate tests to identify changes that may be the result of natural selection. Regions of the genome that are conserved between species are generally considered to be functional, with purifying selection constraining sequences and resulting in lower substitution rates than expected under conditions of neutrality (Cooper et al., 2005). For example, in protein coding genes, the rate of synonymous substitution is generally much higher than the rate of non-synonymous substitution because the latter are under stronger purifying selection. Furthermore, regions of the genome that exhibit accelerated substitution rates may have undergone positive directional selection or relaxation of purifying selection. Identifying these regions with accelerated substitution rates in a phylogenetic framework can therefore provide insight into the selective pressures acting on them and may enable the identification of potential changes in function in lineages of interest (Sackton et al., 2019; Espindola-Hernandez et al., 2022; Kowalczyk et al., 2020; Pollard et al., 2006).

A number of sophisticated methods exist to model how substitution rates in protein-coding genes vary across codons and lineages, such as PAML’s (Yang et al., 1997) branch-site model (Zhang et al., 2005), and models implemented in HyPhy (Pond and Muse, 2005) including aBSREL (Smith et al., 2015) and BUSTED (Murrell et al., 2015), among others. These models have been modified to account for multinucleotide mutations (Lucaci et al., 2021; Venkat et al., 2018), and some have been implemented to estimate changes in selective constraint (e.g. RELAX (Wertheim et al., 2015)). However, comparative studies frequently estimate that, among the 3-8% of vertebrate genomes that are conserved between species, a majority of these regions are non-coding (Siepel et al., 2005; Consortium, 2020). While a number of popular methods exist to estimate simple models of variable conservation and acceleration in non-coding regions of the genome (e.g., PHAST (Siepel et al., 2005; Hubisz et al., 2011), phyloP (Pollard et al., 2010), GERP (Cooper et al., 2005)), these approaches have largely focused on finding regions of conservation amongst the vast quantity of neutrally evolving non-coding regions of the genome. There are thus far few tests that allow researchers to ask whether non-coding regions of the genome are accelerated specifically on branches of interest that may be associated with a trait or trait value of interest.

Of these methods, phyloP (Pollard et al., 2010) from the PHAST (Hubisz et al., 2011) package conducts likelihood ratio tests to identify conservation, acceleration, or substitution rate shifts in a set of pre-specified lineages, modeling substitution rates on the target lineages using a scaling factor relative to the background rate. The BEAST package (Drummond and Suchard, 2010) assumes a random local clock model. They use an indicator variable to denote rate changes in each node and put a Possion prior to control the total number of rate changes on the tree. CoEvol (Lartillot and Poujol, 2011) jointly models genomic substitution rates and continuous phenotypic traits using a multivariate Brownian diffusion process. In the “Forward Genomics” framework (Hiller et al., 2012; Prudent et al., 2016), genome sequences are imputed in ancestral species and compared among species groups with and without the trait of interest to identify associations between presence-absence of genomic loci and phenotypic variation. O’Connor and Mundy (2009) and O’Connor and Mundy (2013) use the likelihood ratio test to detect associations between genotypes and a discrete phenotype. Under the null model (genotype and phenotype are independent), the rate matrices of the genotype and phenotype are independent, while a scaling factor depending on the phenotype is multiplied to the rate matrix of the genotype under the alternative model. TraitRate (Mayrose and Otto, 2011; Levy Karin et al., 2017) also use likelihood methods to detect molecular rate changes associated with discrete phenotypes. While TraitRate and CoEvol model both substitution rates and phenotypic traits of interest, most other approaches use a phylogeny to estimate lineage-specific or sub-tree-specific substitution rates, which are then tested for associations with phenotypic traits of interest. Kowalczyk et al. (2019) developed RERconverge to estimate lineage-specific substitution rates on a phylogeny and demonstrated its use in linking substitution rates and mammalian lifespan (Kowalczyk et al., 2020). However, many of these methods lack complexity compared to their counterparts designed for protein-coding regions, which limit their ability to detect complex patterns of rate shifts, particularly when the species of interest do not form a monophyletic clade.

Recently, we developed PhyloAcc (Hu et al., 2019) (pronounced “Phylo-A-see-see”), a Bayesian method to quantify multiple shifts in substitution rate on a phylogeny. It infers the most probable pattern of shifts in substitution rate from sequence alignments and identifies genomic elements with lineage-specific accelerations using Bayes factors. This opens up the possibility to form hypotheses on links between accelerated rates of substitution on multiple lineages and other traits of interest. For example, PhyloAcc and RERconverge have both been applied to test for correlations between convergent phenotypic states in a phylogeny and substitution rates (Sackton et al., 2019; Hu et al., 2019; Partha et al., 2017; Chikina et al., 2016; Tong et al., 2022). Whereas RERconverge is designed to test one pattern of rate shifts at a time on the tree, PhyloAcc can fit an unrestrained, full model to the input sequences, with rates and rate shifts estimated for each DNA element on each branch of the tree. Such a model allows researchers to ask general questions about genome-wide rate shifts on a phylogeny and their association with phenotypic states, making possible tests for general patterns of evolution (e.g.“Which elements are accelerated on a pre-specified branch or set of branches”; “Which branches have an excess of rate shifts across all elements?”).

Although the methods mentioned above all estimate substitution rates along a phylogeny in different ways to assess shifts in evolutionary rates, they all accept as input a single species tree, and tacitly assume that the gene tree toplogies for all regions of the genome are identical to each other and to the species tree. However, phylogenies for different regions of the genome (which we refer to as gene trees by convention, even for non-genic regions of the genome) can differ from the species history and from other genomic regions due to multiple biological processes such as incomplete lineage sorting (ILS) or deep coalescence, which occurs when variation in ancestral species persisted after speciation, as well as introgression, and gene duplication and loss (Maddison, 1997; Edwards, 2009; Avise and Robinson, 2008). Phylogenetic discordance is commonly observed across the tree of life (Jarvis et al., 2014; Pease et al., 2016; Sun et al., 2021; Lopes et al., 2021) and failure to account for it can lead to mis-estimation of substitution rates when sequences from discordant loci are mapped onto the species tree (Mendes and Hahn, 2016) as well as incorrect inference of divergence times (Angelis and Dos Reis, 2015; Jennings and Edwards, 2005). Hahn and Nakhleh (2016) address the importance of considering gene tree topology variation when attempting to correlate substitution rates and phenotypic traits, specifically in the context of convergent evolution. However, even when the gene tree and species tree are topologically identical, the two can still differ in their branch lengths (Edwards, 2009). Recently, the multispecies coalescent ILS-aware software Bayesian Phylogeography and Phylogenetics (BPP) was extended to include relaxed molecular clocks (Rannala and Yang, 2017; Flouri et al., 2022). However, this model estimates overall rates of each branch of the species tree, as opposed to estimating rates of individual loci along each branch of the species tree. Ogilvie et al. (2017) improves the relaxed random clock model by considering the multispecies coalescence for more accurate inference of per-species substitution rates. In general, macroevolutionary models of molecular clocks and substitution rates have yet to embrace the widespread heterogeneity in gene trees found across the Tree of Life, with unknown consequences for molecular dating, PhyloG2P, and other questions in evolutionary biology (Bravo et al., 2019).

To more accurately estimate substitution rates and identify noncoding sequences that may have experienced accelerated evolution on particular lineages of a tree, here we extend the Bayesian model implemented in PhyloAcc to account for phylogenetic (henceforth “gene tree”) discordance. In our new model, named PhyloAcc-GT, we specify a prior distribution for the gene tree of each element according to the multispecies coalescent model (Rannala and Yang, 2003; Rannala et al., 2020). The full likelihood of the observed sequences from extant species and unobserved sequences from extinct species is defined conditioning on the latent gene tree estimated based on DNA substitution models. To sample gene trees from the posterior distribution, we also develop a Markov chain Monte Carlo (MCMC) algorithm (Liu, 2008) using a new Metropolis-Hastings proposal distribution targeting the conditional posterior distribution of the gene tree conditioning on the species tree, sequence alignment and other parameters. We use sub-tree pruning and re-grafting when proposing new gene tree topologies, but carefully select candidate locations when re-grafting the tree to improve sampling efficiency. Through extensive simulations with various acceleration scenarios, we show that PhyloAcc-GT outperforms both PhyloAcc and *BEAST2 (Ogilvie et al., 2017; Heled and Drummond, 2009), another Bayesian method for detecting substitution rate variation while accounting for ILS. We use PhyloAcc-GT to re-analyze two data sets, one consisting of 43 bird species with a focus on convergent loss of flight in ratites (Hu et al., 2019; Sackton et al., 2019) and the other consisting of 62 mammal species with a focus on convergent evolution of traits linked to marine life (Hu et al., 2019). We show that, after accounting for gene tree discordance PhyloAcc-GT is able to distinguish spurious signals of acceleration due to gene tree variation from true rate shifts. Finally, we also greatly improved the usability and efficiency of our software by developing a command-line user interface that facilitates pre- and post-processing analyses and provides adaptive method selection (PhyloAcc versus PhyloAcc-GT) based on site concordance factors (Ané et al., 2007; Minh et al., 2020) in the input alignments.

## Methods

### Bayesian model to estimate substitution rates in the presence of gene tree discordance

For a given sequence alignment of a genomic element, we estimate substitution rates in the presence of gene tree discordance based on an input species tree, hereafter denoted as ***T***, with branch lengths representing the expected number of neutral substitutions per site and coalescent units from which we calculate population size parameter *θ* ≡ 4*Nμ*, where *N* is the effective population size and *μ* is the mutation rate per site per generation. Parameter *θ*, whose estimation will be discussed later in “Estimating population size parameters”, measures the rate of coalescence in a species and is required when applying the multi-species coalescent model.

Let **Θ** = (*θ*_1_, · · ·, *θ_N_*) denote population sizes for all the *N* species on the species tree. A set of target lineages in the phylogeny to test for acceleration can also be provided if known *a priori*. To model patterns of shifts in substitution rate, PhyloAcc-GT follows the original PhyloAcc model and assumes that substitution rates can only take 3 values corresponding to 3 conservation states. We use ***Z*** = (*Z*_1_, · · ·, *Z_N_*) ∈ {0, 1, 2}^*N*^ to represent these latent conservation states for *N* species on the tree. *Z_s_* = 0 is the background state with the background rate *r*_0_ = 1. *Z_s_* = 1, 2 represent the conserved state and the accelerated state, with the corresponding conserved rate *r*_1_ < 1, and accelerated rate *r*_2_ > *r*_1_. In this way, we frame our test for accelerated substitution rates relative to a pre-measured background or neutral rate of substitution across the genome. Rates are inferred under (up to) three models: a null model that restricts all lineages in ***T*** to the background *r*_0_ or conserved rate *r*_1_, a restricted model in which the target lineages, if present, are allowed to evolve at *r*_2_, and a full model in which all lineages can have any of the three *r* values. Conservation states are defined on each branch of the species tree, and the transition between states is assumed to be Markovian with a state transition probability matrix Φ. The genealogical relationships among sequences of an element are modeled by a latent gene tree variable, denoted by ***G***. The prior distribution of a gene tree given the species tree and population sizes is defined according to the standard multi-species coalescent model. We model DNA sequences evolving according to a continuous-time Markov process defined on the gene tree, whereas the substitution rates are determined by the conservation states in each branch of the species tree.

Under the GTR substitution model, substitutions on one branch of the gene tree follow a continuous-time Markov process with the stationary distribution ***π*** and a rate matrix *Q*. Instead of assuming a fixed and known stationary distribution of the base frequencies, ***π*** = (*π_A_, π_C_, π_G_, π_T_*), for all elements as in the original PhyloAcc, in PhyloAcc-GT we model the stationary distribution of each element independently. Here we use the strand-symmetry model (Bielawski and Gold, 2002; Singh et al., 2009) and assume that substitution rates are the same on the two DNA strands, i.e.,*π_A_* = *π_T_* and *π_G_* = *π_C_*. Thus, we have only one free parameter *π_A_*, for which we impose a half-Beta prior: 2*π_A_* ~ Beta(*γ, γ*). The strand-symmetry assumption can be relaxed, in which case the Beta prior can be replaced by a Dirichlet distribution that can model a vector of probabilities of any finite dimension.

For one element of length *l*, let 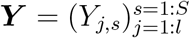 denote the observed aligned sequences in extant species. We use ***X*** = {***Y***, ***H***} to represent the complete data, where ***H*** stands for the unobserved sequences in ancestral species at both coalescent events on the gene tree and speciation events on the species tree. The posterior distribution of all the latent variables (***G***, ***Z***, ***H***) and unknown parameters (***r***, ***π***, **Φ**) is proportional to the product of the likelihood of the complete data given the latent gene tree ***G***, conservation states ***Z***, and parameters ***r***, ***π***, **Φ**, and their joint prior distribution The inference is made by MCMC sampling from this posterior distribution. More details are given in the Appendix D.

### MCMC procedure for posterior inference

Here, inferring the substitution rate ***r*** and the conservation states ***Z*** for each lineage are of the greatest interest, allowing us to identify the most probable pattern of substitution rate shifts along the phylogeny for each element. However, other variables, e.g., the gene tree ***G*** and the ancestral sequences ***H***, cannot be easily integrated out. As such, we use collapsed Gibbs sampling (Liu, 1994) to make posterior inference of all parameters. For each element, we iteratively impute ancestral DNA sequences ***H*** and, conditional on the imputed ***H***, sample conservation states ***Z***, substitution rates ***r***, the stationary distribution of base frequencies ***π***, gene trees ***G***, and the hyper-parameters from their conditional posterior distributions.

We use the forward-backward (Felsenstein, 1973) algorithm to compute conditional likelihoods and sample ***Z*** and ***H***, and use the Metropolis Hastings (MH) algorithm to sample ***r***. Because the substitution rate matrix *Q* depends on *π_A_*, we employ the MH algorithm to sample the posterior distribution of ***π***.

When proposing a new gene tree ***G*** for a given element, we use two MH moves. The first move proposes to change the tree topology of the element. We randomly select a gene tree branch *s*, disconnect the sub-tree rooted at *s* from the remaining tree, and graft it back at a new position in the remaining tree. When proposing the new position, we use the imputed ancestral sequences ***H*** to compute transition probabilities of the sequence from all candidate grandparent nodes compatible with the species tree and the current gene tree structure to *s*. A candidate node is chosen with probability proportional to its transition probability. Such a proposed move takes into account both the sequence information and the tree structure. Second, we update gene tree branch lengths locally by shifting the height of each internal node in the gene tree without altering the gene tree topology using a MH algorithm with uniform proposals centering around the current node position. The correctness of the MCMC algorithm is shown in Appendix B.

The strategy of subtree pruning and re-grafting for updating the tree topology has been explored previously (Rannala and Yang, 2017, 2003). However, to our best knowledge, our design is the first to utilise sequence information to guide the MCMC move directly. Rannala and Yang (2003) randomly select a feasible branch to graft back to, while Rannala and Yang (2017) prefer smaller topological changes by selecting a new position with probability inversely proportional to the number of nodes on the path to the dissolved branch.

### Detecting and Reconstructing Patterns of Acceleration based on Bayes Factors and Estimated Conservation States

PhyloAcc-GT fits up to three nested models to each input alignment and selects the best one based on marginal likelihoods (Bayes factors) of the models.

When a set of target species are specified, we run all three models. Under the null model 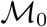, we assume no species is in the accelerated state. Under the lineage-specific model 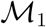, we only allow specified target lineages to be in the accelerated state. Finally, we run a full model, 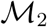, allowing all species not in the outgroup to be in the accelerated state. We identify target lineage-specific accelerations from elements that best fit 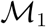 based on two Bayes factors: 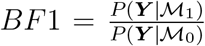, which reflects support for the target-restricted model compared to the conserved model, and 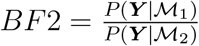, which reflects support for the target-restricted model compared to the unrestricted model. Elements with *BF*1 and *BF*2 greater than some pre-specified thresholds larger than 1 favor the lineage specific model 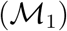, and are most likely to have experienced target lineage-specific accelerations.

PhyloAcc was originally designed to identify convergent rate shifts related to phenotypic convergence, under which it was proven to outperform existing methods. Under such scenarios, target lineages consist of all extant species having the convergent phenotype. However, PhyloAcc can be used more generally, and allows users to specify any combination of lineages as the target set and identify elements that are accelerated within target lineages, or to provide no target lineages to see which elements are best explained by 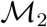. In our application here, as previously (Hu et al., 2019), we do so while also satisfying the condition of Dollo irreversibility of acceleration, although this assumption can be relaxed.

The identified elements that favor 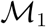 can have varying patterns of acceleration, because not all species in the target group are necessarily accelerated. We identify accelerated lineages by filtering out 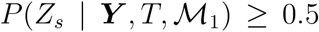 or higher for each lineage *s* in the target group inferred under 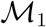. Patterns of acceleration can be similarly inferred based on 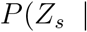 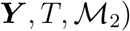 for elements favoring 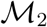 with or without an input target set.

When a target set is not specified, we recommend running both model 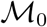 and 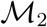 to detect elements experiencing rate acceleration in any lineage. Elements having 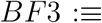 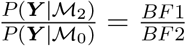 greater than some threshold (at least 1) are likely to have experienced accelerations in some branches of the tree. The precise pattern of acceleration can be inferred from the ***Z*** vector estimated under 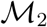, in the same way as under 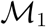, and they imply potential commonalities among accelerated lineages that may not have previously been evident.

To compute the marginal distribution of the observed sequences, we need to integrate out both the gene tree topology and the branch lengths. To do this, we use the Wang-Landau mixture method in Dai and Liu (2020) to estimate marginal likelihoods of the three models, which are in turn used to calculate the Bayes factors. This method works well for both continuous and discrete latent variables. We partition ***Y*** into equally sized data blocks, ***Y***^1^, · · ·, ***Y***^*b*^, and recursively apply the Wang-Landau mixture method with a sequence of target and surrogate distributions. In the first step, we take the prior distribution as the surrogate distribution and 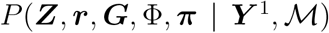 as the target distribution to estimate 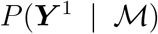. In subsequent step *i*, the target distribution from the previous step 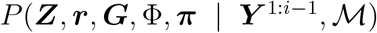 becomes the new surrogate distribution and 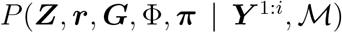 becomes the new target distribution. In the last step, we get an estimate of 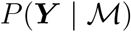.

### Estimating population size parameters

PhyloAcc-GT requires an estimate of the population size, *θ*, which can be challenging in many cases. Some approaches (Flouri et al., 2018; Rannala and Yang, 2017) provide direct estimates of *θ* for current (when more than one allele per extant species is sampled) and ancestral species; other approaches, such as the “two-step” species tree methods, which are helpful in cases of large, genome-wide data sets, estimate branch lengths in coalescent units (t/N), from which *θ* could be extracted if one knows the number of generations per branch (Liu et al., 2015, 2010; Degnan and Rosenberg, 2009; Mirarab et al., 2014). Additionally, whereas some phylogeographic approaches for estimating ancestral population sizes can benefit from the information from multiple loci (Flouri et al., 2018), here we try to estimate rate parameters for a single locus, which alone cannot yield robust estimates of branch-specific population sizes. In our approach, we estimate *θ* first, then treat *θ* as a fixed input that we condition on to estimate other parameters.

For a given branch in a tree, PhyloAcc-GT requires a length *l*_1_ in units of expected number of substitutions per site. This is a common output of phylogenetic software packages (e.g. RAxML (Stamatakis, 2014), IQ-TREE (Nguyen et al., 2015)) and, if estimated from unconstrained sites, can be related to the neutral substitution rate as *l*_1_ = *tμ* where *t* is the number of generations. Other software such as MP-EST (Liu et al., 2010) and ASTRAL (Mirarab et al., 2014)) estimate branch lengths in coalescent units, which are defined with respect to the number of generations *t*. For a given branch, the length in coalescent units is *l*_2_ = *t*/(2*N*). Using these two definitions of branch length, we estimate *θ* on branch *l* as: 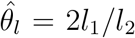. For all extant species, *θ* is set to 0 as only one sequence per extant species is usually available, and *θ* for the root node is set as the average *θ* values among the internal branches of the species tree. PhyloAcc-GT performs this calculation internally both with the species tree provided by the user, with branch lengths in units of expected substitutions per site under the neutral rate, as well as with a topologically identical species tree with branch lengths in coalescent units estimated using one of the methods mentioned above. If this second tree is not pre-estimated, PhyloAcc-GT automates its estimation with a Snakemake pipeline that uses IQ-TREE to estimate individual locus trees for up to 5,000 of the longest input loci and ASTRAL to obtain branch lengths in coalescent units.

### Simulating sequence data

To test the accuracy of PhyloAcc-GT and compare it to other methods, we simulated sequence data given a species tree under several scenarios of substitution rate acceleration, where we allow either a single monophyletic acceleration, two independently accelerated clades, or three independently accelerated clades (Fig. 2). We simulated sequences using the “SIMULATE” function in PhyloAcc-GT. The SIMULATE function takes as input a species tree with branch lengths in expected number of substitutions, population size parameters, a DNA substitution stationary distribution, and a rate matrix *Q*. For each element, the function first generates a gene tree according to the multi-species coalescent model (Appendix C), and the DNA sequence at the root of the gene tree following a simulated stationary distribution based on the Beta distribution: 2*π_A_* ~ *Beta*(10, 10). Subsequent sequences are generated using the continuous time Markov model, but only those for extant species are output. The conserved and accelerated rates are generated from Gamma distributions: *Gamma*(5, 0.04) and *Gamma*(10, 0.2), respectively. The two distributions correspond to a mean rate of 0.2 and 2. The population size parameters for the simulations are estimated from real data based on ratites (see below). For our simulations, we first simulated 400 loci with conserved rates in every lineage. Then, for each scenario outlined above, we combined these 400 loci with up to 100 loci simulated with accelerated substitution rates in the specified lineages. All elements are simulated to be 100 base pairs (bp) long.

We used these simulated datasets in several ways to compare PhyloAcc-GT’s accuracy in identifying both genomic elements experiencing acceleration and lineages harboring those elements that are accelerated. First, we calculated the area under the precision-recall curve (AUPRC) based on BF1. Precision is the proportion of true positives out of all called positives. Recall is the percentage of true positives identified out of all positives. When a dataset contains many more negatives (i.e., elements without any acceleration along the tree) than positives (i.e., elements having at least one acceleration event on a target lineage), the precision-recall curve has been shown to be a more informative measure of a method’s performance than receiver operating characteristic (ROC) curves (Davis and Goadrich, 2006). AUPRC varies as a function of the proportion of positives in the dataset (Saito and Rehmsmeier, 2015), measuring model performance under different degrees of data skewness. We therefore vary the ratio of the number of accelerated to the number of conserved conserved elements from 1 to 100, and compare AUPRC between PhyloAcc-GT and the original PhyloAcc species tree model (henceforth just “PhyloAcc”).

We also examined how well PhyloAcc-GT identifies specific lineages with accelerated substitution rates under the optimal model inferred. Here we compared the performance of PhyloAcc-GT, PhyloAcc, and the random local clock model implemented in *BEAST2 (Ogilvie et al., 2017). *BEAST2 also estimates substitution rates along a phylogeny within a Bayesian framework, but does not restrict rate variation to three distinct classes. Because *BEAST2 does not explicitly calculate the probability of acceleration per lineage for a given element, to compare the performance of *BEAST2 with that of PhyloAcc-GT and PhyloAcc, we estimate *P* (*Z* = 2 | ***Y***) by the proportion of MCMC outputs in which the branch is accelerated. We treat a branch to be in the accelerated state if its estimated rate is greater than the estimated rate of its parent branch. *BEAST2 does not require input *θ*, but models and integrates out population size. However, for a fair comparison with PhyloAcc-GT, we input and fix the theta parameters to *BEAST2 as well.

To test how PhyloAcc-GT and PhyloAcc handle phylogenetic discordance, we adjusted *θ* in our simulated data. When *θ* increases, the mean and variance of coalescent times between sister lineages on the tree increase, leading to an increased probability of discordance. We multiplied the *θ* values estimated from the ratite data by 3, 6 or 10 and use these new parameters to simulate new sequences under the three previously described scenarios.

### Ratite and marine mammal data

To further compare PhyloAcc-GT with PhyloAcc, we use data from two systems: birds and mammals. We previously analyzed these data with PhyloAcc and identified genomic elements associated with loss of flight in birds (ratites) and the transition to aquatic lifestyles in mammals (marine mammals) (Hu et al., 2019). The bird dataset consists of 43 species, including 9 flightless birds (ratites: ostrich, moa, 2 species of rhea, emu, cassowary, and 3 species of kiwi), 27 volant bird species, and 7 reptiles as outgroup species A.1. We used the alignment of 284,001 conserved non-coding elements, the species tree, and genome-wide estimates of neutral substitution rates from Sackton et al. (2019) and Hu et al. (2019).

For the mammal data, we previously used (Hu et al., 2019) alignments of 283,369 conserved non-coding elements from 62 species, a species tree A.2, and genome-wide estimates of neutral substitution rates from the UCSC 100-way vertebrate alignment (Blanchette et al., 2004). We identified conserved non-coding elements using PHAST (Hubisz et al., 2011) and estimated neutral substitution rates from 4-fold degenerate sites using phyloFit (Hubisz et al., 2011); see Sackton et al. (2019) and Hu et al. (2019) for full description of these methods. From these datasets, since we are interested in comparisons of PhyloAcc-GT with PhyloAcc, we limit our comparisons to the elements previously inferred to be accelerated in either ratites (806 elements based on Bayes factor cutoffs of log *BF*_1_ > 20 and log *BF*_2_ > 0) or marine mammals (2,106 elements based on Bayes factor cutoffs of log *BF*_1_ > 5 and log *BF*_2_ > 5) (Hu et al., 2019).

For both datasets, we estimate **Θ** based on the species tree topology as described above, using gene trees from 20,000 randomly selected loci. For each set of gene trees, we ran MP-EST five times and used the branch lengths from the run with the maximum likelihood. 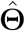 is then calculate based on the branch lengths two trees (one with branch lengths in units of relative number of substitutions and one with branch lengths in coalescent units) as outlined in the section above (“Estimating population size parameters”). We repeated this process 50 times and averaged the *θ*s as the population size parameters for each dataset. We used the estimates from the ratite data as 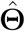 for the simulated data sets described above.

### Site concordance factors

To reduce run time, we use site concordance factors (sCF) to determine on an element-by-element basis whether to use the PhyloAcc-GT method, which accounts for phylogenetic discordance in the input locus, or the original PhyloAcc species tree method, which uses only a single species tree for all elements. Concordance factors (Ané et al., 2007; Baum, 2007) were first implemented on a per-site basis by Minh et al. (2020) in IQ-TREE2 (Minh et al., 2020) to summarize discordance among genes relative to a species tree. Briefly, sCF is calculated for a given branch in the species tree by first calculating concordance factors among sub-alignments of quartets of species sampled from that branch (*CF_q_*). For each quartet we count the number of sites in the alignment of those species that match the topology in the species tree (e.g. ((A,A),(G,G)) and divide that number by the total number of decisive alignment sites (see Minh et al. (2020), Equation 2). In IQ-TREE-2 (Minh et al., 2020), these values of *CF_q_* are calculated over all sites in every input alignment and averaged to obtain an overall summary of discordance in the dataset. Here, we re-implement the sCF calculation to be applied to each individual locus, resulting in a value for each branch in the species tree for each locus. We then use the sCF values for each locus to guide the selection of the PhyloAcc gene tree or species tree method. This can be specified in two ways by the user: 1) if the average of all sCF values for the locus are below some threshold this locus will be run with the gene tree method, otherwise it will be run with the species tree method, and 2) if the proportion of branches with a sCF below some threshold exceeds another threshold, this locus will be run with the gene tree method, otherwise it will be run with the species tree method. Thresholds are specified with user inputs and are meant to limit the number of loci run with the computationally more intensive gene tree method.

### Benchmarking with simulated data

We benchmarked both the PhyloAcc-GT and PhyloAcc species tree algorithms by using simulated datasets. We simulated loci on species trees of various sizes (9, 13, or 17 species). For each species tree, we simulated 100 sequences of various length (100, 200, 400, and 600bp) and ran each locus through both programs in batches of 10 loci with each batch using 4 threads. We measured average run time and average maximum memory use on each batch and divided by batch size to get average resource use per element. We ran these benchmarks on the Harvard Research Computing Cannon Cluster.

## Results

The PhyloAcc-GT algorithm is implemented in a C++ codebase that accounts for phylogenetic discordance in the input loci while estimating substitution rates across a phylogeny. This algorithm, along with the original PhyloAcc codebase, which uses a single species tree for all input loci, and a newly implemented command-line user interface, are packaged together to form the PhyloAcc software (https://phyloacc.github.io/). The user interface is implemented in Python and provides the ability to easily batch input elements into separate runs for PhyloAcc, which can be partitioned between the species tree or gene tree methods. These batches are then executed via an automatically generated Snakemake file that can submit batches in parallel as separate jobs to a high-performance computing cluster with job scheduling software (e.g. SLURM).

### Model performance with correct input targets

To measure their ability to differentiate accelerated elements from non-accelerated ones with respect to a set of target lineages, we input the correct (i.e. simulated) target set to PhyloAcc-GT and PhyloAcc, using three sets of simulated data (single accelerated clade; two independent accelerations; and three independent accelerations; see Fig. 2). We then measure the area under precision-recall curve (AUPRC) of logBF1 while varying the proportion of accelerated elements. We find that PhyloAcc-GT has high precision and recall as measured by AUPRC. As the proportion of target-specific accelerated elements decreases, it becomes harder to detect these elements from the remaining conserved ones because more conserved elements can be falsely identified as accelerated at any fixed logBF1 cutoff. However, the AUPRC for PhyloAcc-GT never falls below 95% regardless of the type of acceleration scenario or the fraction of input elements that are accelerated (Fig. 3). By contrast, the original PhyloAcc always has a lower AUPRC, especially when the truly accelerated lineages are a sub-set of the input targets (e.g. Fig. 3D). When the ratio of conserved to accelerated loci is 100:1, PhyloAcc-GT can identify true positive cases more than 95% of time, while PhyloAcc’s performance can drop to 75%. The precision-recall curves at ratio 50:1 conserved to accelerated loci are also shown in Fig 3.

In addition to assessing model selection accuracy by element, we also check for accuracy of predicting lineages with accelerations by examining the posterior probability of rate acceleration in each branch *P* (*Z* = 2|***Y***) under the most favored models based on Bayes Factors. We find that both PhyloAcc-GT and PhyloAcc can precisely identify accelerations occurring on the terminal branches of the species tree. However, PhyloAcc-GT is much better at identifying accelerations on internal branches of the tree than PhyloAcc (Fig. 4). Under the multispecies coalescent, gene tree branch lengths for extant species are longer than the branches of the species tree, whereas the same is not necessarily true for internal branches (Fig. 1). As such, PhyloAcc tends to overestimate substitution rates along terminal branches more than along internal branches.

**Figure 1:**
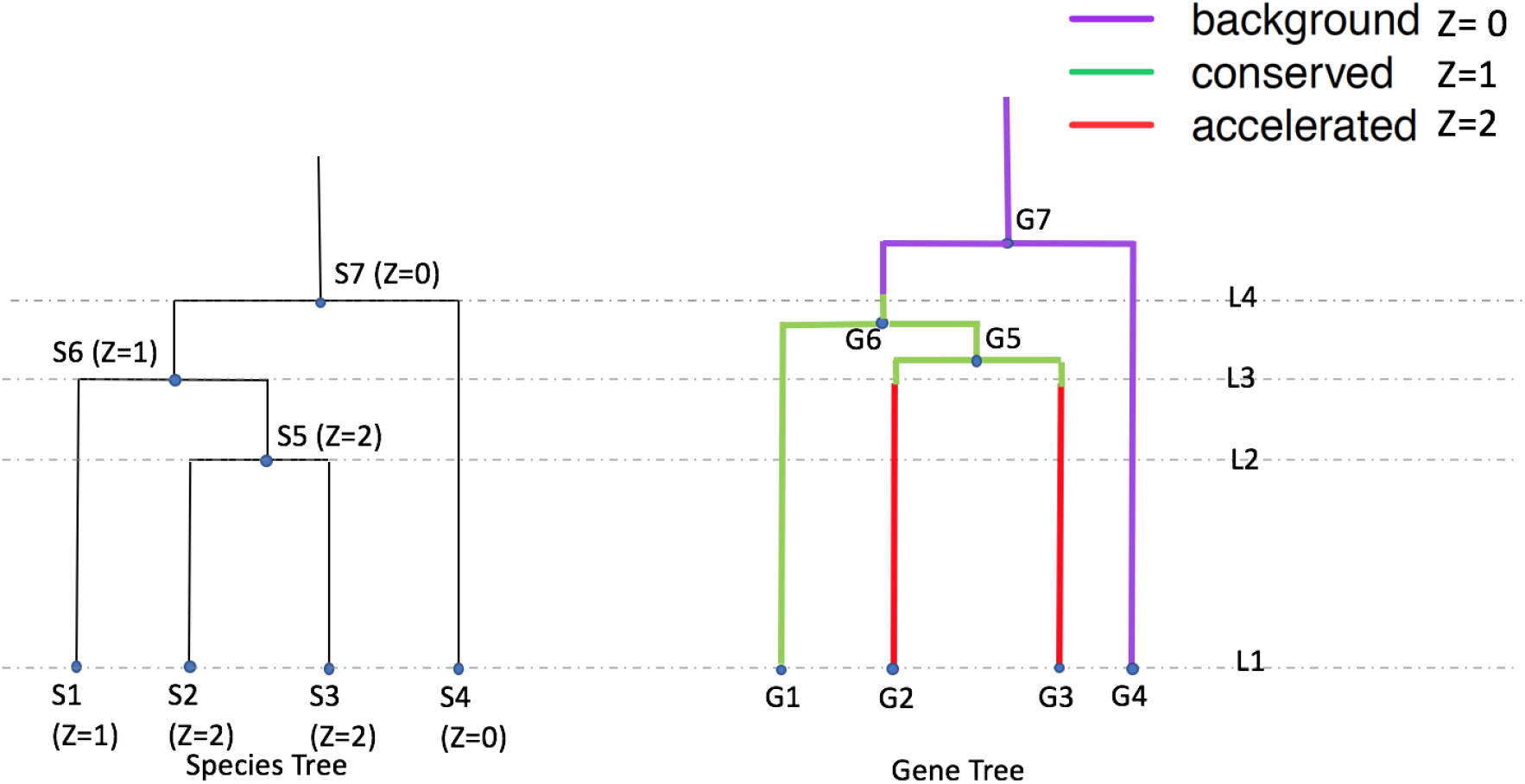
Conservation states and DNA evolution given a species tree and a gene tree. On the left is a species tree with given rate conservation states at each node (Z=0, 1, or 2). On the right is a gene tree for a single input element. The sequences of the element evolve according to the gene tree. In different species tree branches, substitution rates are different and determined by ***Z***. A gene tree branch crossing multiple species can be in different conservation states indicated by different colors in the figure.

We also compare the ability of PhyloAcc-GT to detect accelerated lineages to *BEAST2. We find that *BEAST2 reports lower posterior probabilities for accelerated lineages for most branches than both PhyloAcc-GT and PhyloAcc (Fig. 4 A, C, and E). The average estimated posterior probabilities for acceleration across accelerated branches are 0.62 for the single acceleration case, 0.59 for two accelerated clades, and 0.5 for three accelerated clades. These values, while generally over 0.5, fall below a conservative threshold that one may use to identify accelerated lineages. Additionally, *BEAST2 has less resolution in discerning accelerated lineages from non-accelerated lineages, with several non-accelerated lineages having an average posterior probability of acceleration above 0.5, which may lead to a higher false positive rate in detecting accelerated elements on a given branch (Fig. 4 B, D, and F).

### Model Performance with mis-specified targets

To test the ability of PhyloAcc-GT to distinguish target-specific acceleration from off-target acceleration using logBF2, we consider three scenarios where the specified target lineages include only some or none of the simulated accelerated lineages (Fig. 5). In scenario (1), the input target species partially overlap truly accelerated species: we simulate two independently accelerated clades, and specify one of them as the target lineage and the other as a non-accelerated clade. In scenario (2), the input target species are a subset of the truly accelerated species: we simulate three independently accelerated clades, and specify as targets only one of those clades. In scenario (3), the truly accelerated species do not intersect with input target species. Area under the ROC curve (AUROC) between PhyloAcc and PhyloAcc-GT are recorded in Fig 5 legend. We use AUROC to measure model performance because the input set of targets and specified set of targets can be any two acceleration patterns. It is reasonable to not assume that elements accelerated under one pattern (the input target set under model 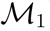) are significantly more frequent than the other (the input target set under Model 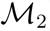). Both methods are highly accurate in excluding non-specific accelerated elements. AUROC are close to 1 as presented in Table 1. We also compute the true positive rate (TPR) at 1% and 5% false positive rate cutoffs. In all scenarios, PhyloAcc-GT has higher accuracy than PhyloAcc.

**Table 1:** Comparing true positive rate at different false positive rate cutoffs using logBF2 to distinguish target-specific accelerated elements from non target-specific accelerated elements under different scenarios of target mis-specification between PhyloAcc-GT and PhyloAcc. Truly accelerated species either overlap (rows 1 & 2), include (row 3 & 4) or are completely different from input target species (rows 5 & 6).

| Testing Scenario | Method | TPR@1%FPR | TPR@5%FPR |
| --- | --- | --- | --- |
| 1 | PhyloAcc-GT | 0.89 | 0.96 |
|  | PhyloAcc | 0.76 | 0.84 |
| 2 | PhyloAcc-GT | 0.99 | 1 |
|  | PhyloAcc | 0.92 | 1 |
| 3 | PhyloAcc-GT | 0.97 | 0.98 |
|  | PhyloAcc | 0.94 | 0.97 |

Next we assess the inference of conservation states, specifically *P* (*Z* = 2|***Y***), or the probability of acceleration along a given branch, of all branches by PhyloAcc-GT and PhyloAcc under the above scenarios of target mis-specification. Results using *BEAST2 are not presented because it does not allow prior selection of targets.

We find that PhyloAcc-GT is more accurate in identifying accelerated branches than PhyloAcc (Fig. 6). Although PhyloAcc-GT produces a slightly wider range of acceleration probabilities across truly accelerated lineages than PhyloAcc, almost all probabilities are still above 0.75. Consistent with the previous analysis, PhyloAcc-GT performs much better than PhyloAcc in detecting accelerations along internal branches. For non-accelerated branches, both methods tend to have higher estimated posterior probabilities of acceleration in clade C compared to other non-accelerated species (e.g., scenario 1). The higher probabilities are probably due to the shorter branch lengths of C1 and C2, and their proximity to truly accelerated branches. Compared with the case of a single acceleration in Figure 4 when 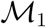 is the true model, correctly identifying 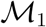 in PhyloAcc-GT or PhyloAcc can reduce the posterior probability of acceleration in non-accelerated branches. However, as these posterior probabilities are still below 0.5 in most elements, the ability in inferring the correct acceleration pattern and the number of independent acceleration events is largely not affected by the input target species.

### Identifying accelerated lineages with no input target set

Although a model that tests for accelerations on specific target lineages may prove a better fit than a full model, often this information is unavailable, or we may want to ask general questions about our sample (e.g. “How many elements show acceleration in any lineage?", “Which lineages have the most accelerated elements?”). To test PhyloAcc-GT’s performance under such scenarios, we use the same simulation setting as previously (Fig. 2) but now use logBF3 to identify accelerated elements, and then *P* (*Z* = 2 | ***Y***) under model 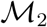 to reconstruct the patterns of acceleration.

**Figure 2:**
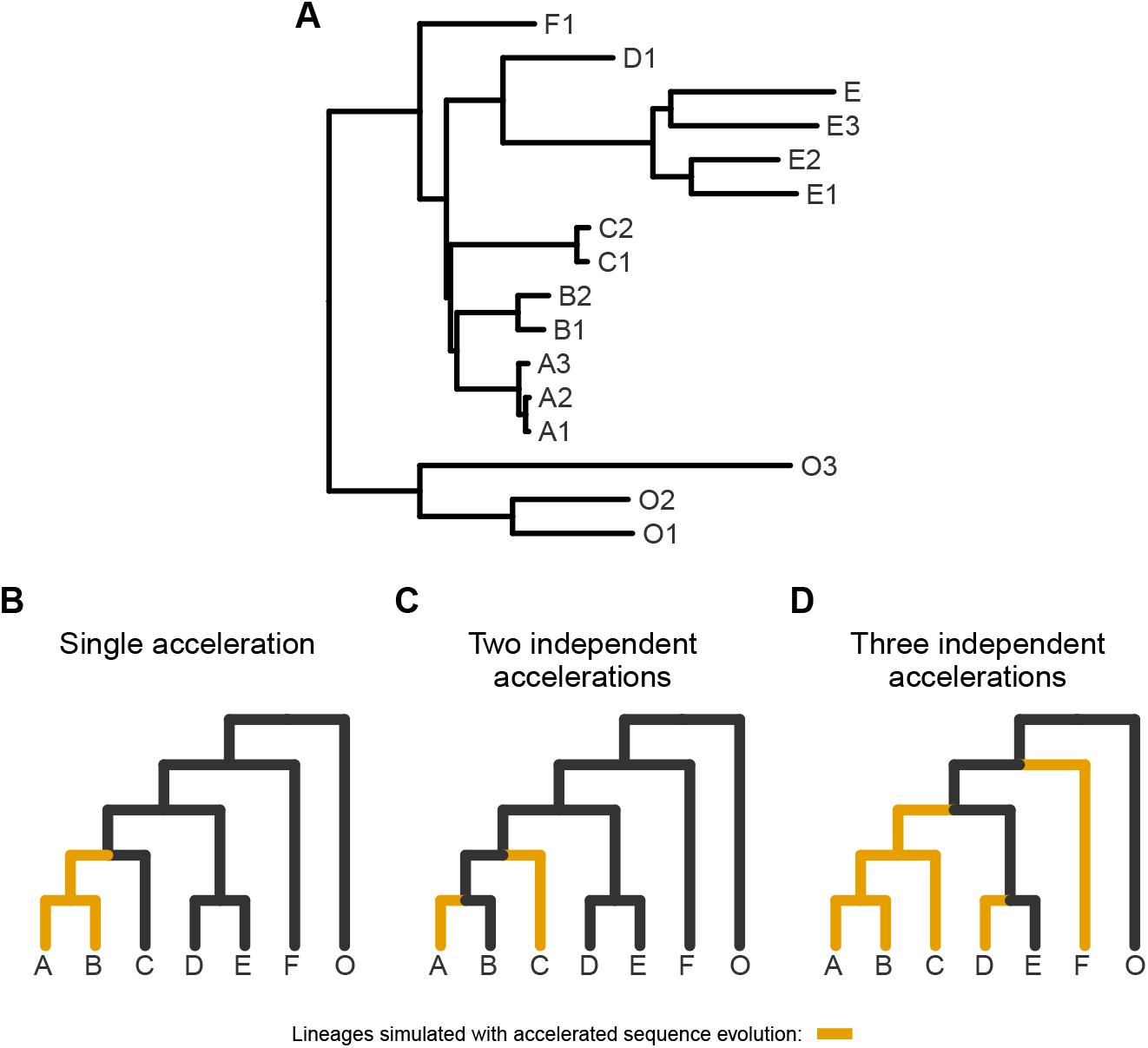
Trees representing simulated scenarios of accelerated sequence evolution. A) The full tree used for simulations with topology and branch lengths based on the ratite phylogeny (Fig. A.1). For visualization only, B-D represent collapsed versions of the tree in A with arbitrary branch lengths and tip labels representing monophyletic clades. A) A single monophyletic acceleration. B) Two independent accelerations. D) Three independent accelerations.

We again find that PhyloAcc-GT more accurately identifies accelerated elements than PhyloAcc in all scenarios (Fig 7). The differences in performance by the two methods are more pronounced as the percentage of non-accelerated elements in the data increases, and the performance gap is larger than when testing a set of target lineages with logBF1 (Fig 3). We also find similar patterns in the distribution of *P* (*Z* = 2 | ***Y***) for truly accelerated branches whether we input the correct target set or not (Fig 8 v.s., Fig 4). However, when identifying accelerated lineages for a given element without specifying targets, we see larger variation in *P* (*Z* = 2 | ***Y***) among non-accelerated branches that are nearby truly accelerated branches on the species tree (Fig 8 B, D, and F, compared to the results when target branches are specified (Fig 4 B, D, and F), and truly accelerated branches with short branch lengths (e.g., clade A). However, these posterior probabilities generally do not exceed 0.5 for non-accelerated branches, and are mostly above 0.5 for truly accelerated branches. When only a single clade is truly accelerated, we observe more variation in posterior probabilities when an input set is not specified. In this case, when accelerated lineages are correctly specified in the input set, no false positives are observed among 17 non-accelerated branches under 100 simulations. When using 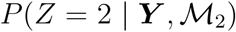, the false positive rate is 4% and the false negative rate increases from 3% to 9%.

**Figure 3:**
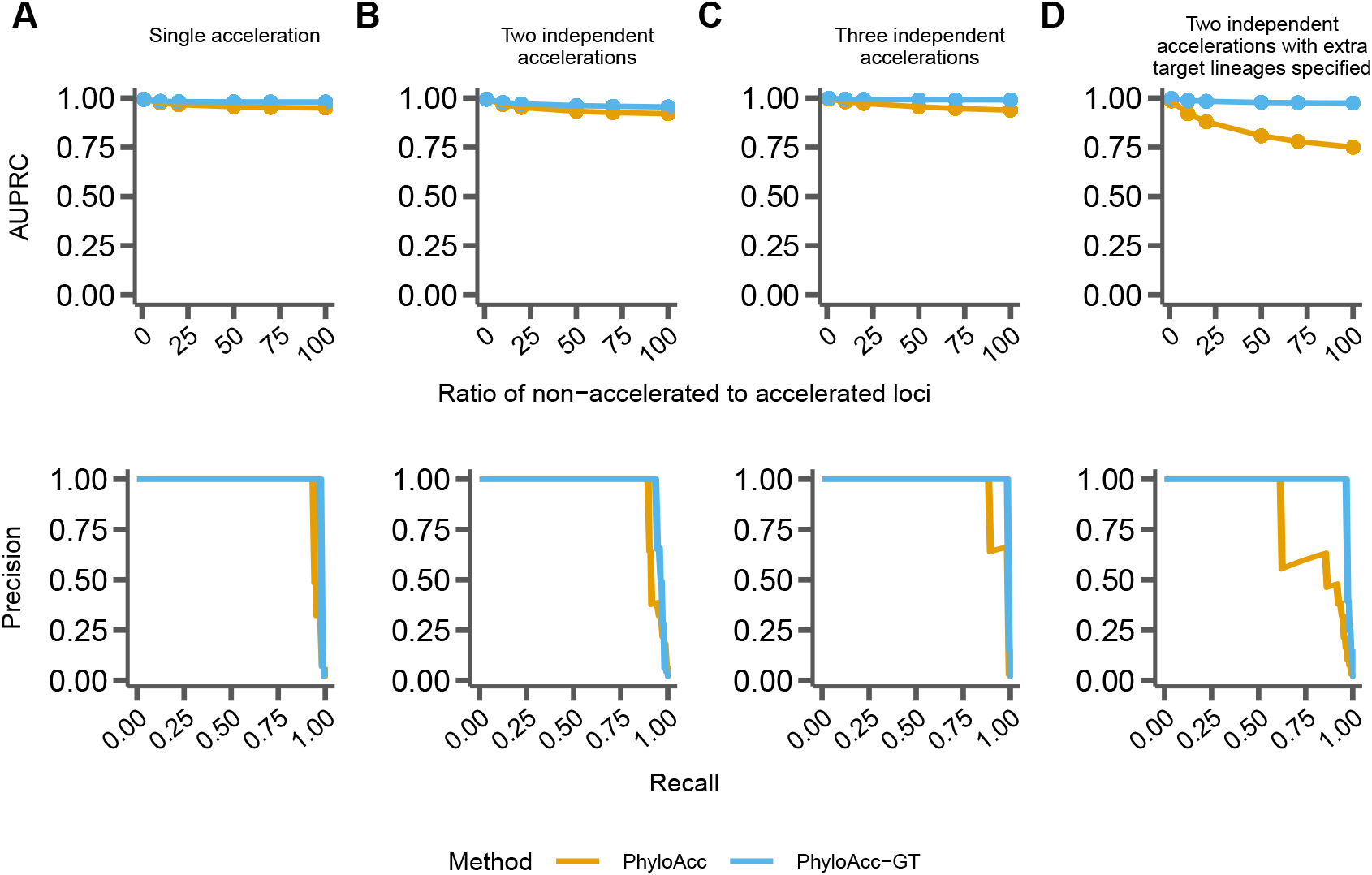
Comparing performance between PhyloAcc (orange) and PhyloAcc-GT (blue). The top row shows the Area Under the Precision-Recall Curve (AUPRC) while varying the ratio of simulated conserved to accelerated loci. The bottom row shows a single precision recall curve at a ratio of 50 conserved loci per accelerated locus. In A-C, the specified target lineages match those lineages on which accelerations were simulated. A) Loci simulated with a single monophyletic acceleration. B) Loci simulated with two independently accelerated clades. C) Loci simulated with three independently accelerated clades. D) Loci simulated with two independently accelerated clades, but with additional target lineages provided to each method.

**Figure 4:**
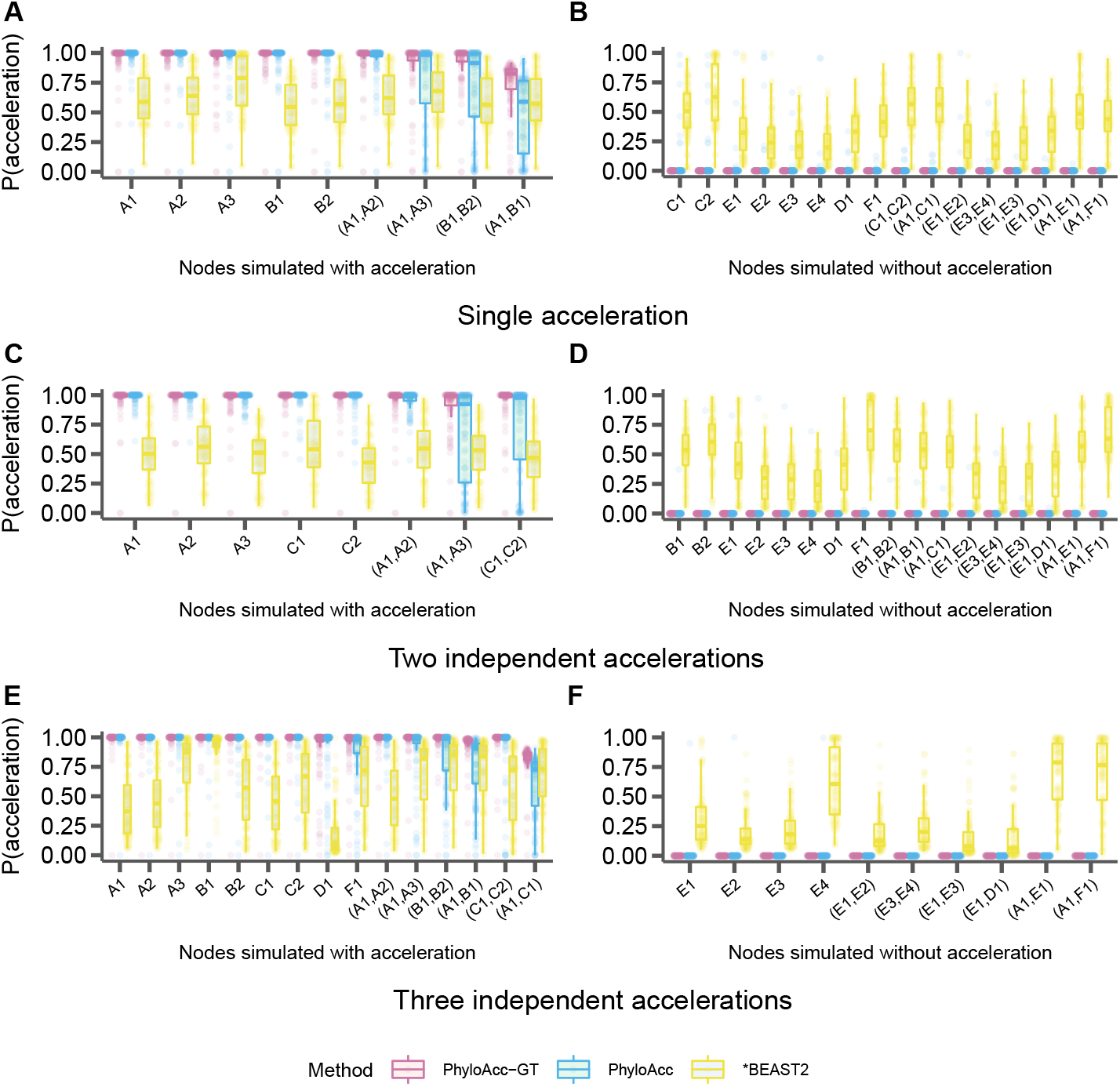
Comparison of the identification of lineage specific rate accelerations between three methods, PhyloAcc-GT (purple), PhyloAcc (blue), and *BEAST2 (yellow) when the input target lineages match truly accelerated lineages. Each distribution corresponds to the estimated *P* (*Z* = 2 | ***Y***)s of a branch from 100 simulated elements. Branches are indicated on the x-axis of each plot and correspond to those in Fig. 2A. Distributions on the left correspond to lineages simulated to have accelerated sequence evolution in each of the three scenarios in Fig. 2, whereas distributions on the right correspond to those without accelerated sequence evolution. A & B: The probability of acceleration for each locus and lineage using sequences simulated with a single accelerated clade (Fig. 2B). C & D: Probability of acceleration for each locus and lineage using sequences simulated with two independently accelerated clades (Fig. 2C). E & F: Probability of acceleration for each locus and lineage using sequences simulated with three independently accelerated clades (Fig. 2D).

**Figure 5:**
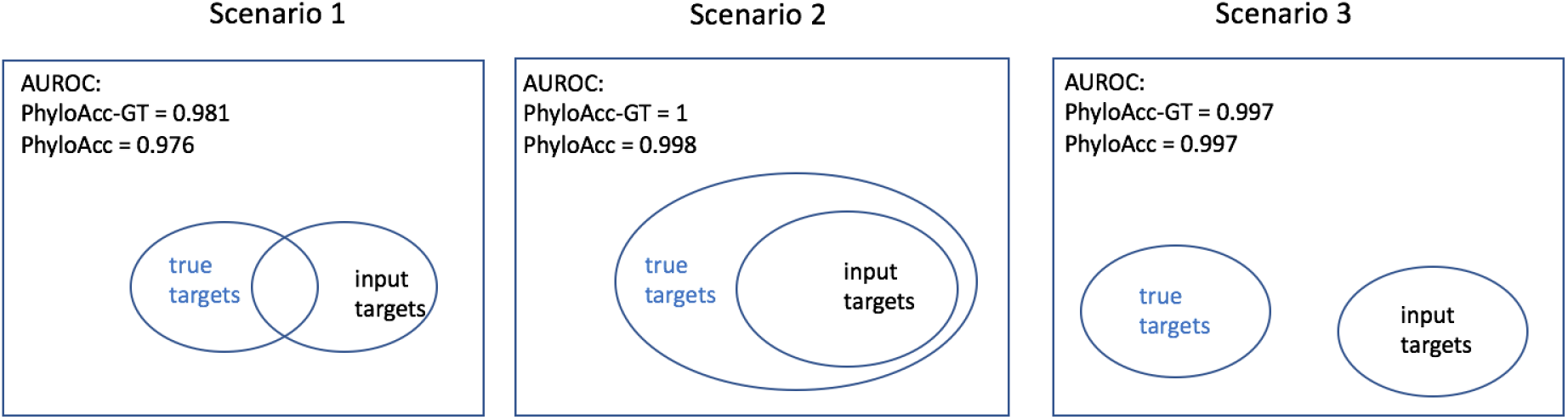
Scenarios for testing model performance with mis-specified targets, along with AUROC for both PhyloAcc-GT and PhyloAcc.

**Figure 6:**
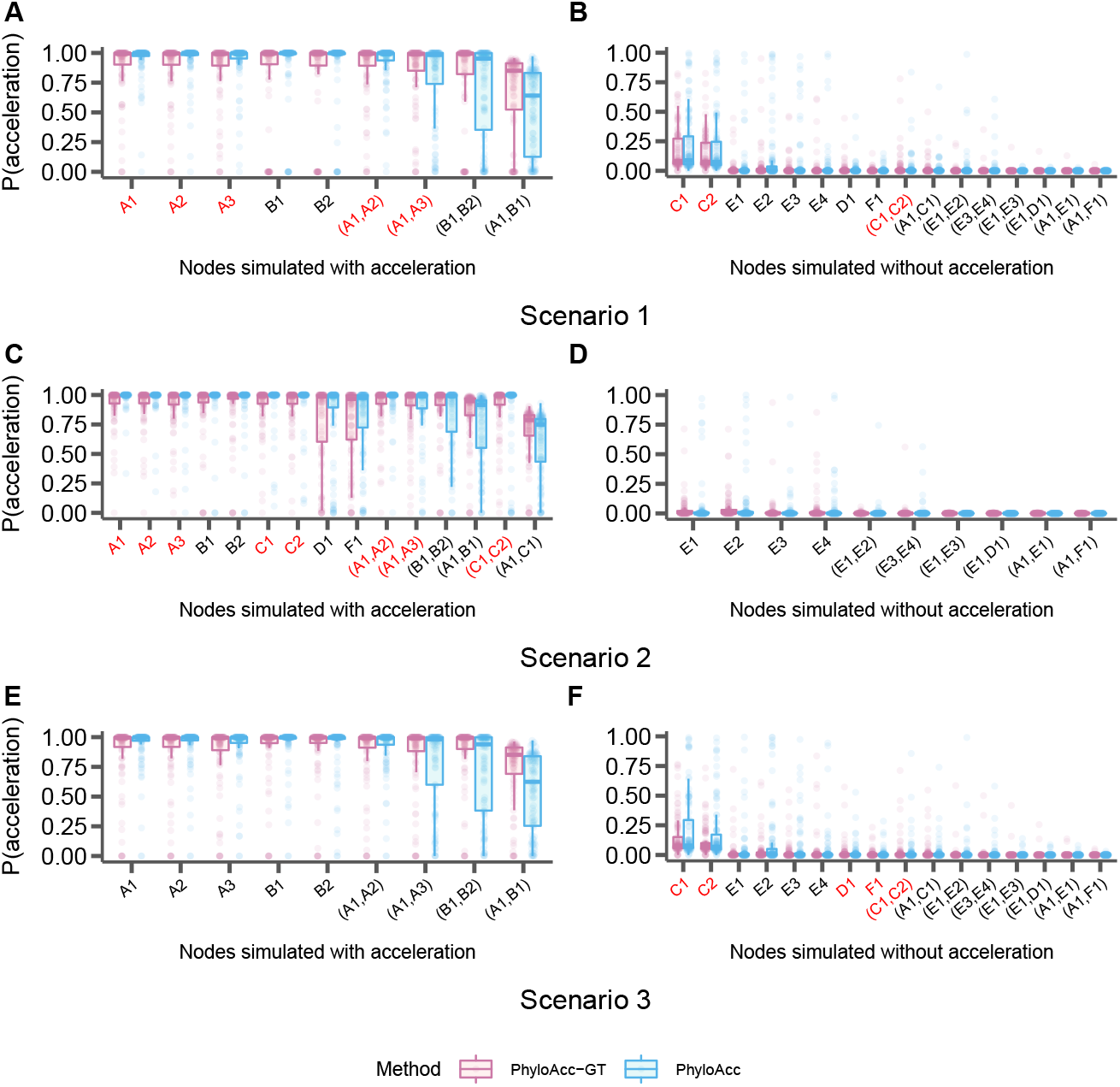
Distributions of the probability of acceleration (*P* (*Z* = 2 | ***Y***)) for each branch in the input species tree when specified target lineages are mis-specified. Branches are indicated on the x-axis of each panel and correspond to those in Fig. 2A. Distributions on the left correspond to lineages simulated to have accelerated sequence evolution in each of the three scenarios in Fig. 2, and distributions on the right correspond to those without accelerated sequence evolution. Branches in red on the x-axis are those that were specified as target lineages for M1 in each run of PhyloAcc or PhyloAcc-GT and the three scenarios correspond to those outlined in Fig. 5. Each point represents one simulated locus. A & B: The probability of acceleration using sequences simulated with a single monophyletic acceleration (Fig. 2B) and targets specified that partially overlap the truly accelerated lineages. C & D: The probability of acceleration using sequences simulated with two independent accelerations (Fig. 2C) and targets specified as a subset of the truly accelerated lineages. E & F: The probability of acceleration using sequences simulated with three independent accelerations (Fig. 2D), and no truly accelerated lineages specified as targets.

**Figure 7:**
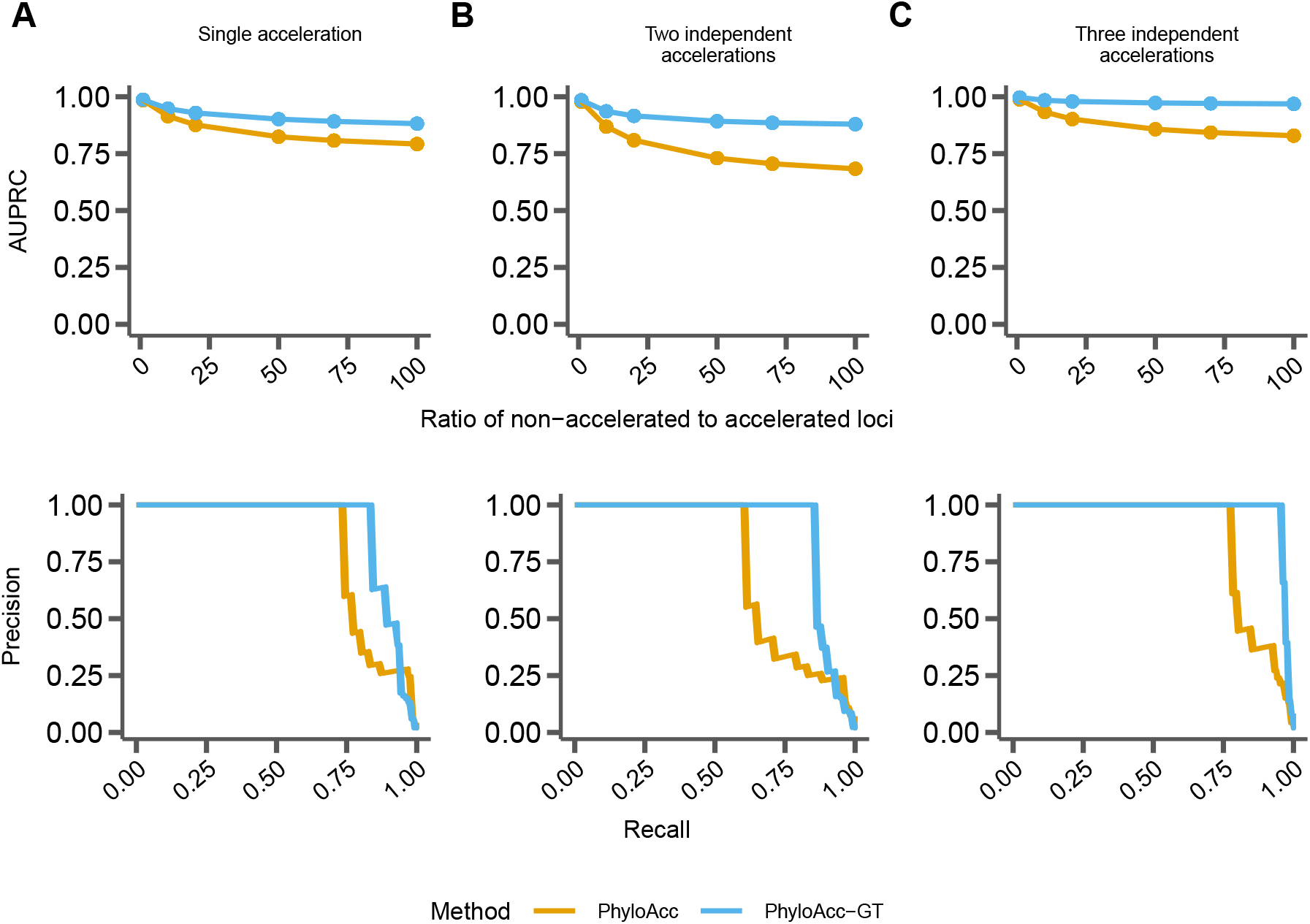
Comparing performance between PhyloAcc (orange) and PhyloAcc-GT (blue) without specifying target lineages. The top row shows AUPRC for both methods while varying the ratio of non-accelerated to accelerated loci. The bottom row shows a single precision-recall curve at a ratio of 50 non-accelerated loci per accelerated locus. A) Loci simulated with a single, monophyletic acceleration. B) Loci simulated with two independent accelerations. C) Loci simulated with three independent accelerations.

**Figure 8:**
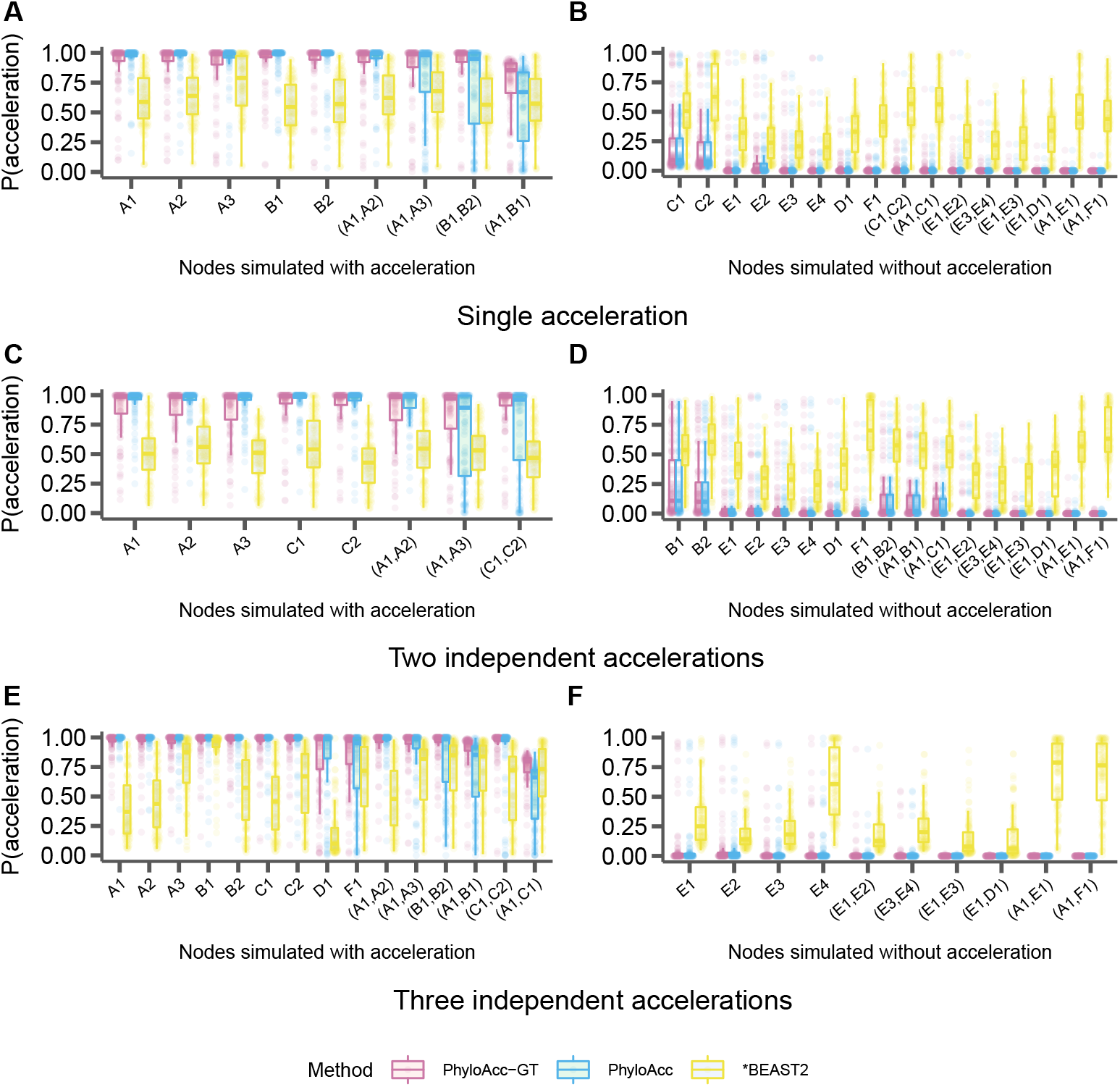
Comparison of the identification of lineage specific rate accelerations between three methods, PhyloAcc-GT, PhyloAcc, and *BEAST2, when no target lineages are provided (i.e. from 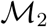). Each distribution corresponds to the estimated *P* (*Z* = 2 | ***Y***)s of a branch from 100 simulated elements. Branches are indicated on the x-axis of each plot and correspond to those in Fig. 2A. Distributions on the left correspond to lineages simulated to have accelerated sequence evolution in each of the three scenarios in Fig. 2, whereas distributions on the right correspond to lineages simulated without accelerated sequence evolution. A & B: The probability of acceleration for each locus and lineage using sequences simulated with a single accelerated clade (Fig. 2B). C& D: Probability of acceleration for each locus and lineage using sequences simulated with two independent accelerations (Fig. 2C). E & F: Probability of acceleration for each locus and lineage using sequences simulated with three independent accelerations (Fig. 2D).

This result implies that specifying a target set is beneficial, and if one has logical target lineages in mind, we recommend using them to reconstruct patterns of acceleration using results from 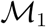 for those selected elements, to achieve a slightly lower false positive rate. However, if an input set cannot be specified, our method still reliably identifies accelerated elements and infers patterns of acceleration using 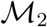, with only minor reductions in accuracy.

### Robustness to phylogenetic discordance

The amount of phylogenetic discordance present within the input loci affects the identification of both elements and lineages experiencing accelerated substitution rates. To assess how PhyloAcc-GT performs with varying levels of phylogenetic discordance due to ILS, we varied the population size parameter *θ* in each of our three simulation cases. We find that in each case when considering logBF1, as *θ* increases the AUPRC of PhyloAcc-GT decreases depending on the fraction of elements that are truly accelerated (Fig. 9). However, in every case PhyloAcc-GT achieves a higher AUPRC than PhyloAcc, especially when the *θ*’s are large and the proportion of accelerated elements is low.

**Figure 9:**
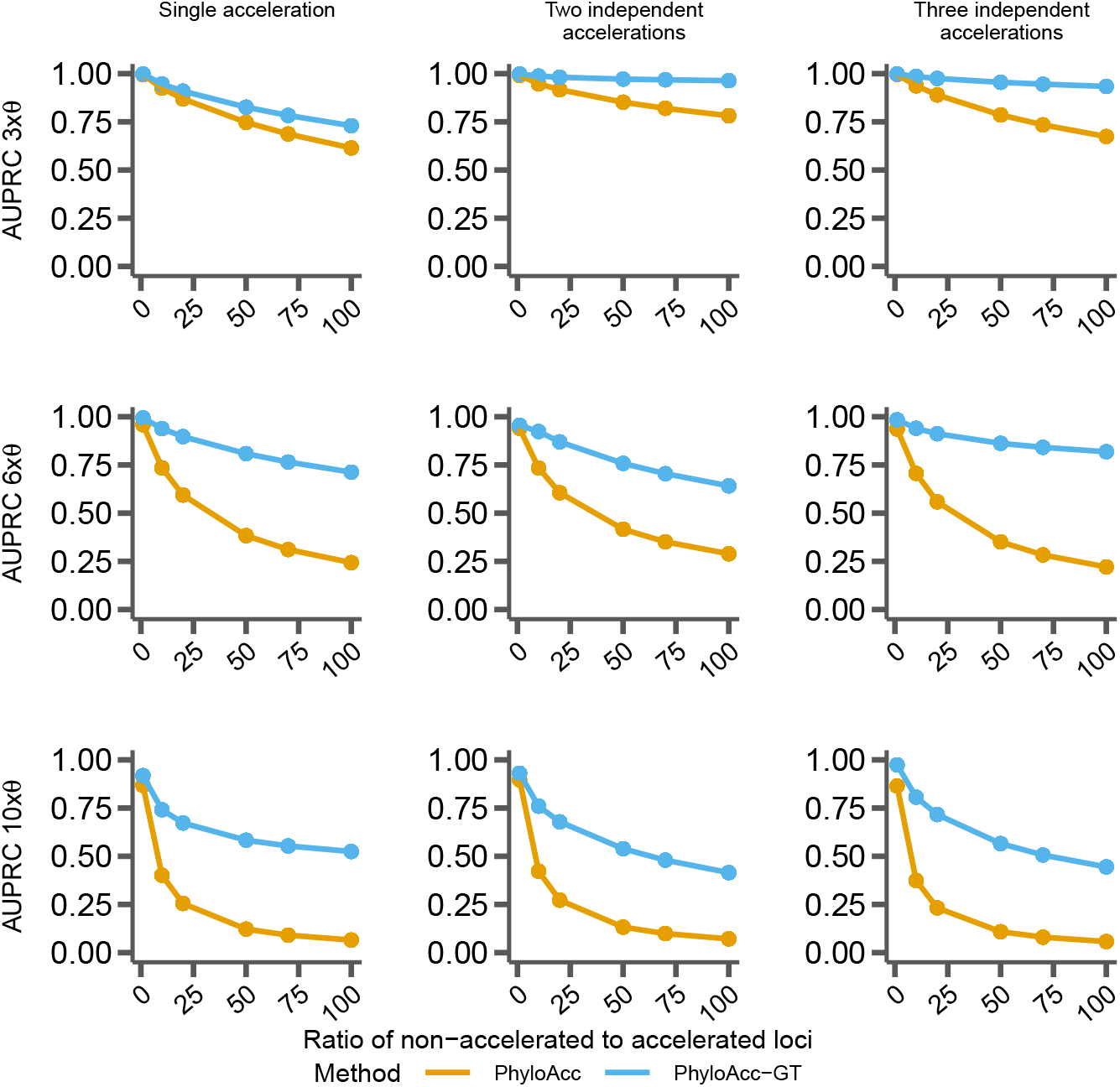
Comparing performance between PhyloAcc (orange) and PhyloAcc-GT (blue) with varying levels of **Θ** and the ratios of non-accelerated to accelerated loci. Rows represent different scales of the *θ* (3x, 6x, or 10x) values estimated from the ratite dataset (see Methods) while columns represent different simulation scenarios.

We also find that PhyloAcc-GT consistently outperforms PhyloAcc in identifying accelerated lineages while minimizing false positives, regardless of the extent of ILS (Fig.10, H.1 and H.2). For PhyloAcc-GT, the posterior probabilities for accelerated branches are mostly above 0.75 and in most cases close to 1, while the probabilities are close to 0 for non-accelerated branches. Again, we see that PhyloAcc also performs quite well when identifying accelerations on terminal branches of the species tree, but its performance on internal branches is greatly affected by the amount of ILS. In many cases, the average posterior probability for acceleration on a truly accelerated internal branch falls below 0.2 and even close to 0 for very high levels of ILS. In general, *BEAST2’s performance does not seem to be affected by varying amounts of ILS. Accelerated lineages also consistently have an average probability of acceleration > 0.5 when analyzed with *BEAST2. However, in most instances this probability is less than 0.75 and has large variation. *BEAST2 also has a high variance in posterior probabilities for non-accelerated branches, which are routinely between 0.25-0.5, and can be up to 0.75 in some branches, possibly leading to false positives.

**Figure 10:**
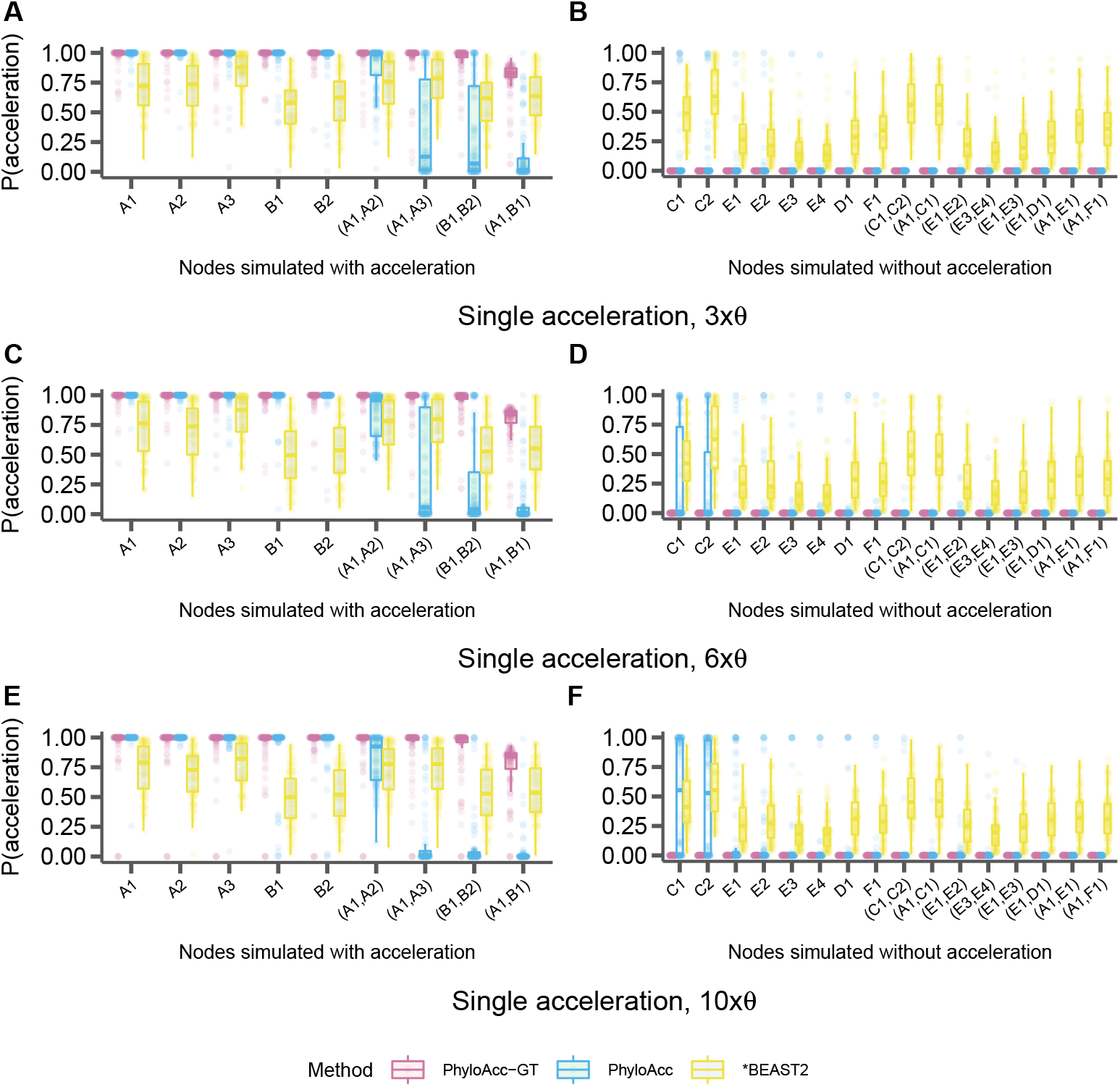
Distributions of the probability of acceleration (*P* (*Z* = 2 | ***Y***)) using PhyloAcc-GT, PhyloAcc and *BEAST2 while scaling the population size parameter, **Θ**. Distributions shown for data simulated with a single acceleration in the A and B clades (Fig. 2B). Branches are indicated on the x-axis of each panel and correspond to those in Fig. 2A. Distributions on the left correspond to lineages simulated to have accelerated sequence evolution while the distributions on the right correspond to lineages simulated without accelerated sequence evolution. A & B: Sequences simulated with 3 times the expected ***θ***. C & D: Sequences simulated with 6 times the expected ***θ***. E & F: Sequences simulated with 10 times the expected ***θ***.

### Robustness to mis-specification of theta

Because *θ* is a parameter calculated by our model and is key to determining the amount of phylogenetic discordance in the input data, we test the performance of PhyloAcc-GT when it is mis-specified. Under mis-specification of *θ*, we still identify numerous elements that favor a model of target-specific acceleration with both BF1 and BF2 being positive. We find that PhyloAcc-GT correctly identifies accelerated elements over 97% of the time for when the scaling factor of *θ* is between 0.5 to 2 (our tested cases). At 5% FPR, the true positive rates are all above 0.98 across scenarios (Table 2).

**Table 2:**
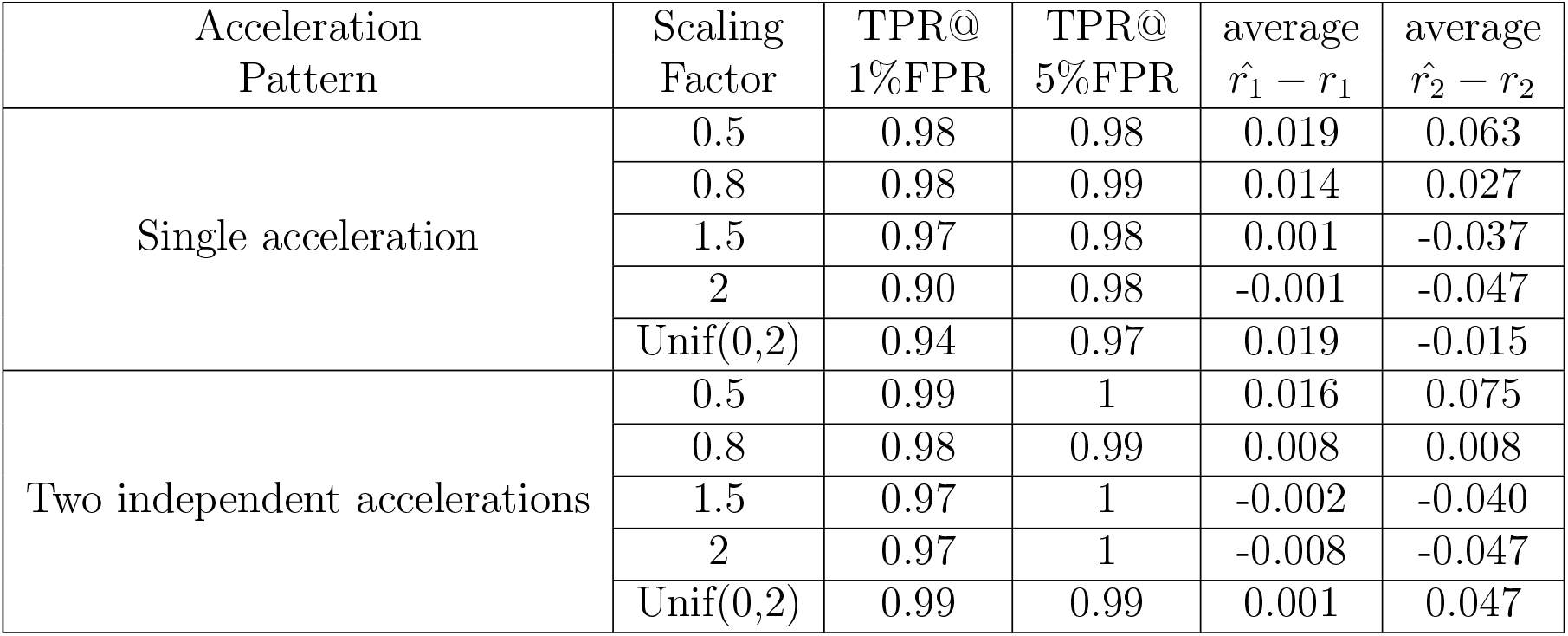
Test sensitivity of PhyloAcc-GT to *θ* mis-specifications. TPRs at the 2 FPR cutoffs are computed based on logBF1 among null elements and accelerated elements.

| Acceleration<br>Pattern | Scaling<br>Factor | TPR@<br>1%FPR | TPR@<br>5%FPR | average<br>$\hat{r}_1 - r_1$ | average<br>$\hat{r}_2 - r_2$ |
| --- | --- | --- | --- | --- | --- |
| Single acceleration | 0.5 | 0.98 | 0.98 | 0.019 | 0.063 |
|  | 0.8 | 0.98 | 0.99 | 0.014 | 0.027 |
|  | 1.5 | 0.97 | 0.98 | 0.001 | -0.037 |
|  | 2 | 0.90 | 0.98 | -0.001 | -0.047 |
|  | Unif(0,2) | 0.94 | 0.97 | 0.019 | -0.015 |
| Two independent accelerations | 0.5 | 0.99 | 1 | 0.016 | 0.075 |
|  | 0.8 | 0.98 | 0.99 | 0.008 | 0.008 |
|  | 1.5 | 0.97 | 1 | -0.002 | -0.040 |
|  | 2 | 0.97 | 1 | -0.008 | -0.047 |
|  | Unif(0,2) | 0.99 | 0.99 | 0.001 | 0.047 |

In addition to model selection, estimates of the conserved and accelerated substitution rates, *r*_1_ and *r*_2_ respectively, are influenced by *θ* as well, though in general the biases tend to be small. When we specify underestimated *θ*s, the model will overestimate *r*_1_ and *r*_2_ and vice versa. When each branch’s input *θ* value is a random scaling of the true *θ*, the direction of estimated bias depends on all the realized *θ*’s along the tree (Table 2).

### Identifying accelerated elements in ratites

We apply PhyloAcc-GT to the 806 conserved non-coding elements previously detected by PhyloAcc (Hu et al. (2019)) to have strong evidence for ratite-specific acceleration (BF1>20 and BF2>0), possibly linking them to the loss of flight. When accounting for phylogenetic discordance with PhyloAcc-GT, we find that marginal likelihoods imply that 88% (713) of the elements still favor *M*_1_, indicating ratite-specific acceleration, whereas 8% (67) of those elements previously identified now fall under *M*_0_ and do not show any rate acceleration. Examining the 67 elements favoring *M*_0_, we found that 11 of these elements do not have any target lineages with a high probability to be in the accelerated state (P(Z=2|Y > 0.5) under PhyloAcc (Appendix G).

To determine which elements still show strong evidence of ratite-specific accelerations after accounting for phylogenetic discordance with PhyloAcc-GT, we first determined new Bayes factor cutoffs for the ratite data based on simulated data. We find that the ratio of BF1 between PhyloAcc and PhyloAcc-GT for data generated under *M*_1_ (two accelerated clades) is 1.8, meaning BF1 tends to be higher when using PhyloAcc. To account for this, we adjust our BF1 cutoff to identify ratite-specific accelerations when using PhyloAcc-GT from 20 down to 10. The BF2 cutoff remains 0. Using these cutoffs, we identify 509 out of the original 806 elements (63%) with strong evidence for ratite-specific acceleration. The average estimated accelerated rate (r2) is 2.5, while the mean conserved rate (r1) is 0.16. 88% of these elements have accelerated rate greater than 1, and 56% are greater than 2. Similar to PhyloAcc’s result, the rhea clade is most likely (60%) to experience acceleration among all lineages. Almost all accelerations in this clade are inferred to have occurred in the most recent common ancestor of the two extant rhea species, rather than two independent accelerations. The emu and cassowary branches are the second most likely (40%) lineages to be accelerated, and 80% of the accelerations occurred along their ancestral branch. The ostrich branch is the least likely extant species to have experienced accelerations.

Among accelerated elements, 291 are inferred to have accelerated on only one branch by PhyloAcc-GT. 43% of these single-branch accelerations occur along the ancestral rhea branch, followed by 11% in moa and 11% in the most recent common ancestor of cassowary and emu. The original PhyloAcc, without considering ILS, detected only 265 single-branch accelerations. In some cases, PhyloAcc inferred separate accelerations in sister branches, whereas PhyloAcc-GT infers only a single acceleration in the ancestral branch of the two sibling branches. For example, PhyloAcc estimates element mCE1745684 having two independent accelerations in cassowary and emu, whereas PhyloAcc-GT infers the acceleration to have occurred in their parent species.

Recently an alternative but weakly supported species tree for palaeognaths has been advocated, suggesting that rheas are sister to kiwis, emus, cassowaries, and tinamous Simmons et al. (2022). Re-running PhyloAcc using the alternative tree identifies 817 (log-BF1>20, log-BF2>0 as in Hu et al. (2019)) elements being accelerated. Among these elements, 717 elements overlap with the 806 elements (89%) identified using the original tree. For the remaining elements that are detected under the original tree but not in alternative tree, 77 elements still have the maximum marginal likelihood under model 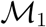, i.e., favoring a pattern of ratite-specific acceleration over no acceleration or acceleration in non-ratites. When running PhyloAcc-GT with the alternative tree, PhyloAcc-GT selects 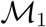 as the optimal model in 713 elements. 671 elements (94%) show evidence of ratite-specific accelerations under both species tree specifications, indicating that PhyloAcc-GT is more robust (the percentage of overlap of estimated accelerated elements is more under PhyloAcc-GT (94%) than under PhyloAcc (89%) to different species tree topologies than PhyloAcc.

### Identifying accelerated elements in marine mammals

We also re-ran PhyloAcc-GT on 1,276 conserved non-coding elements that were previously inferred to have marine mammal specific accelerations using the original PhyloAcc species tree model with BF1 and BF2 cutoffs of 4 (Hu et al., 2019). We find that 1,034 (81%) elements still have the highest marginal likelihood under model 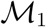, while 225 (17.6%) elements now favor the null model. Setting cutoff at 2 for both log Bayes factors, we estimate 882 elements to have strong target lineage-specific acceleration. The average conserved rate is 0.17 and the average accelerated rate is 2.66, with 761 elements having an accelerated rate greater than 1.

Using PhyloAcc-GT, we find that the branch leading to dolphins experiences the largest number of rate accelerations (606), followed by killer whale (539). Additionally, 403 accelerations occurred in the ancestral cetacean branch. These results differ from using the original PhyloAcc model, which identified, only 279 accelerations in the ancestral cetacean lineage. Among the elements identified as accelerated in this branch by PhyloAcc-GT, PhyloAcc is more likely to identify the acceleration in only one of the two extant species (dolphin or killer whale), with 26 elements actually identified as having independent accelerations in both. For example, for element VCE173687, PhyloAcc estimates a posterior probability of acceleration of 0.89 in the killer whale branch, but only 0.64 in dolphin. However, PhyloAcc-GT infers that there is an acceleration event the ancestral cetatcean branch, and the posterior probabilities of acceleration of the parent and child branches are all greater than 0.88. Other than this difference, inference of conservation states of other target species are the same: both PhyloAcc and PhyloAcc-GT infer an independent acceleration in manatee with posterior probability greater than 0.99, and posterior probabilities of being in the accelerated states for seal and walrus are all below 0.7.

The number of accelerations in manatee, seal, and walrus are 219, 205 and 235, respectively. As opposed to the cetacean clade which has many accelerations in the ancestral branch, in the pinniped clade, most rate shifts happen independently in either the walrus or seal lineages. Only 77 elements are estimated to have experienced one acceleration along the ancestral pinniped branch. This is similar to PhyloAcc’s result: there are 201, 190 and 235 elements accelerated in manatee, seal, and walus, and 65 accelerations in walrus and seal started in their parent species.

### Benchmarking & implementation

We benchmarked PhyloAcc-GT and the original PhyloAcc by running the programs on loci simulated on species trees of various sizes with sequences of varying length. We find that run times for both programs vary depending on both the number of species in the input phylogeny and the length of the input alignment. However, for the gene tree model, sequence length is the more important factor, with simulated data sets with more than 9 species having roughly the same run times, though this is likely dependent on which branches species are added to. We find that for short sequences (100 bp), average run times per element range from 14 to 46 minutes depending on the number of species in the phylogeny (Fig. 11 A). However, as sequence length increases run time also increases substantially. With a sequence length of 400bp, using a tree with 9 species we find an average run time per element of 155 minutes, but using a tree with 13 species average run time per element is on average 460 minutes (Fig. 11 A). For the species tree model, run times are still correlated with both sequence length and tree size, but are substantially reduced compared to the gene tree model. With the species tree model, average times per element range from just 1.5 seconds in a tree with 9 species and elements 100bp long to 17 seconds in a tree with 17 species and sequences 600bp long (Fig. 11 A). The ratite dataset contains 284,001 non-coding DNA elements with a median length of only 103bp, meaning that real datasets should be mostly confined to these lower run time estimates (Fig. 11 B). Memory use also scales with tree size and sequence length, but always remains below 200MB.

**Figure 11:**
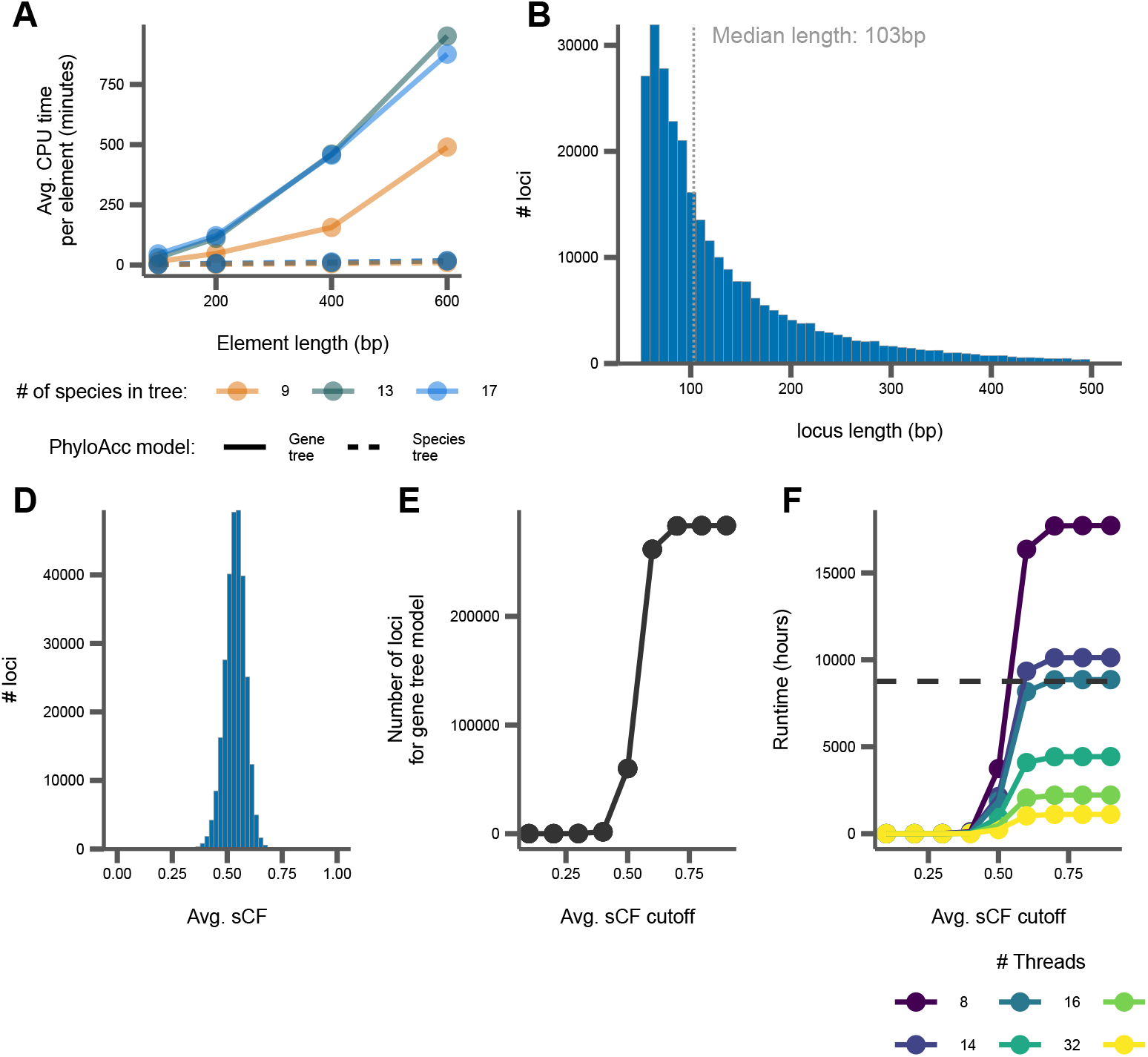
Summaries of benchmarking and concordance factor analysis. A) Average CPU time per simulated element in minutes. B) Distribution of element lengths from ratite data, with the median length labeled and indicated by the dotted grey line. C) Distribution of sCF per element from ratite data. D) The number of ratite elements that would be run with the gene tree model with various sCF cutoffs. E) The expected run time for all elements to complete with the gene tree model from the ratite dataset with various sCF cutoffs. The colored lines correspond to varying the number of threads per element. The dashed black line corresponds to a time of 1 year.

As these benchmarks show, the sampling of locus trees implemented in the gene tree model is a computationally intensive process, requiring substantial CPU resources and time to infer substitution rates even for a single element compared to the species tree model. To address this, we have implemented an adaptive model selection procedure in the user interface that uses site concordance factors (sCF) calculated on each locus to determine whether or not they need to be run with the computationally intensive PhyloAcc-GT, or if the original species tree model in PhyloAcc will suffice. Users provide cut-off values to determine which loci will be run through which model. We show that for the ratite dataset, the average sCF per locus is above 0.5, meaning for most elements, more than 50% of sites support the relationships inferred in the species tree (Fig. 11 C). We varied the average sCF cutoff for these data to see how many loci would be run through PhyloAcc-GT as opposed to the PhyloAcc species tree model and the subsequent effect on estimated run time (assuming linear scaling with increased threads) for the loci that are input to PhyloAcc-GT (Fig. 11 D and E). We find that both the number of loci and the estimated run time both increase as the average sCF cutoff is increased, sometimes becoming excessive with run times over 1 year. However, with a low enough cutoff (e.g. below 0.4), we achieve more reasonable run times when only using PhyloAcc-GT on loci with many discordant sites in many branches of the tree.

With the user interface we also provide summary statistics for the input alignments as well as the option to pre-batch files for submission to a compute cluster via Snakemake. This batching further reduces run time as batches can be run in parallel.

## Discussion

Detecting complex patterns of substitution rate variation in specific lineages of a phylogeny is an important task that may facilitate the association between small-scale sequence evolution with other biological processes, such as structural variation, habitat or environmental shifts, or even phenotypic evolution (Smith et al., 2020; Partha et al., 2019). However, most tests for rate variation across the tree are usually restricted to protein coding regions (Yang et al., 1997; Pond and Muse, 2005) and nearly all such methods for detecting such shifts, whether designed for coding or non-coding regions, do not account for incomplete lineage sorting and deep coalescence, which can arise in many commonly encountered situations and can induce false signatures of rate variation when ignored Mendes and Hahn (2016). Here we present PhyloAcc-GT, which extends PhyloAcc to detect shifts in substitution rate of non-coding elements on phylogenetic trees in the presence of deep coalescence. Through simulation we have shown that accounting for gene tree variation significantly reduces false positive rates when detecting rate acceleration on specific branches. PhyloAcc-GT has higher AUPRC than PhyloAcc, especially when the number of conserved elements significantly outnumbers the number of accelerated elements. PhyloAcc-GT is also superior to PhyloAcc and *BEAST2 in identifying patterns of acceleration along a phylogenetic tree and their associated rates. Compared to *BEAST2, PhyloAcc-GT is more confident in identifying all branches undergoing acceleration, for both tips and internal branches. Compared to PhyloAcc, PhyloAcc-GT has better power in identifying internal branches that are accelerated, resulting in more accurate estimation of substitution rates and inference of whether an element experienced multiple independent accelerations or a single acceleration in an ancestral species. With the introduction of logBF3, which tests support for a model that allows rate acceleration on any lineage, PhyloAcc and PhyloAcc-GT can also be used to test more general hypotheses about molecular evolution in a given phylogeny, such as quantifying which loci are accelerated across the most lineages or which lineages contain the most accelerated elements.

PhyloAcc-GT also provides flexibility in allowing different stationary distributions of DNA substitution models across the genome by inferring the distribution for each element from the data. Simulations (Appendix F) show that modeling the stationary distribution of each element leads to better inference of substitution rates than PhyloAcc, which uses a fixed stationary distribution across all elements and can show poor performance when this global distribution differs significantly from the distribution of a given element. Here we have assumed the strand-symmetry model of DNA substitution *π*; however, the model is easily extendable to other substitution models and priors, such as the Dirichlet distribution. Applying PhyloAcc-GT to accelerated elements in genome-wide bird and mammal data sets, we find that nearly 20% of the elements previously identified by PhyloAcc as accelerated in specific target lineages are likely spurious due to false signatures of acceleration induced by incomplete lineage sorting. Thus, for these two data sets, both of which are known to experience incomplete lineage sorting, PhyloAcc results in substantial improvements in our ability to identify truly accelerated elements.

An important challenge in considering gene tree variation in the PhyloAcc framework is obtaining parameters of population size *θ* for each branch of the species tree. Estimating *θ* for each branch from sequence data or from gene trees is challenging in part because rate variation among loci can mimic variation in coalescence times among loci, sometimes causing identifiability problems (Yang, 1997; Zhu and Yang, 2021). Currently, our approach uses separate estimates of branch lengths in substitutions per site (via concatenation) and in coalescent units (via a species tree method such as MP-EST (Liu et al., 2010) or ASTRAL (Mirarab et al., 2014)) on a pre-specified species tree to obtain estimates of *θ*, which can therefore vary from branch to branch. This approach likely incurs biases, because, even when working with the same species tree topology, the branch lengths obtained via concatenation are likely mis-estimated and do not precisely correspond to branch lengths in a species tree obtained via coalescent approaches (Edwards, 2009; Edwards et al., 2016; Rannala et al., 2020). Additionally, it is well known that methods such as ASTRAL and MP-EST that rely on estimating species tree branch lengths from fixed gene trees estimated in a separate, previous step, result in overestimates of ancestral *θ* (Yang, 1997, 2002). Still, our analysis of the bird and mammal data sets shows that *θ*s obtained in this manner yields reasonable values of *θ*, with small differences in *θ* for most branches, as expected. Additionally, our simulations shows that PhyloAcc-GT is robust to mis-specification of *θ* when model selection is the focus. However, it can overestimate substitution rates when *θ*s are consistently underestimated, and underestimate them when *θ*s are consistently overestimated. When working with data generated from the null model, using underestimated *θ*s leads to PhyloAcc-GT detecting more false positive cases, while using overestimated *θ*s do not seems to result in more false positive. Adjusting the stringency of model selection via the Bayes Factors will be useful in modulating the false positive rate in PhyloAcc-GT.

PhyloAcc and PhyloAcc-GT together provide a flexible framework to identify changes in substitution rates along phylogenetic trees with or without deep coalescence. Our current implementation (https://phyloacc.github.io/) also incorporates many improvements in ease of installation (through bioconda) and use. Although the increased model complexity of the gene tree model (PhyloAcc-GT) provides increased accuracy in the presence of ILS, it also incurs increased use of computational resources, sometimes becoming realistically intractable (Fig. 11). This naturally comes with the additional cost of higher energy use and a larger carbon footprint when running the more complex model, which is becoming an increasing concern for bioinformatics software developers (Grealey et al., 2022). Considering the trade-off between the increased accuracy of a more complex model and the increased resource use those models require, it is valuable to develop novel heuristics to guide users to the appropriate method for the given data – in essence not every locus may need to be analyzed with the most complex model. In our case, we developed an adaptive method selection (PhyloAcc vs. PhyloAcc-GT) for different loci within a data set using site concordance factors (sCF; Minh et al. (2020)) to determine the loci that may be most impacted by phylogenetic discordance. By varying the cut-offs for sCF required to run a locus with the PhyloAcc-GT model, we can drastically reduce run time and energy use with minimal impact on analytical results (Fig. 11), though some post-hoc analyses may be required to assess rates of error.

Going forward, accurate detection of loci across the genome undergoing rate changes in specific target lineages must eventually grapple with well-known complexities of the genome. For example, our current models assume a single neutral set of branch lengths for the species tree for comparison with individual loci. However, different regions of the genome likely experience different neutral substitution rates, thereby requiring greater model complexity (Eyre-Walker and Eyre-Walker, 2014; Hodgkinson and Eyre-Walker, 2011). One way to improve the accuracy of estimation of substitution rates with PhyloAcc might be to use the regions flanking each conserved element to estimate the local neutral substitution rate for a given locus. Additionally, here we have assumed that all branches in the accelerated rate class share a single substitution rate. This constraint can easily be relaxed to allow independent accelerations on a tree to have different rates. As currently implemented, our model assumes the Dollo’s irreversibility condition such that after an acceleration event occurs on a branch for a given element, all descendent species remain in the accelerated state. This assumption could be relaxed by allowing for some probability of reverting from an accelerated to a conserved state via the ***Z*** matrix. PhyloAcc and PhyloAcc-GT currently focus on conserved non-coding elements that use standard models of nucleotide substitution. Arguably, the much large number of conserved non-coding elements than genes or exons in genomes and their likely widespread role in driving phenotypic evolution make a focus on non-coding variation a profitable place to start (Sackton et al., 2019; Lewis et al., 2019; Marcovitz et al., 2016; Mattick, 2005). However, we can extend this model to detect rate shifts in protein-coding regions as well. Finally, for 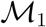, PhyloAcc and PhyloAcc-GT currently focus on sets of target lineages that are in or not in a designated target set or are characterized by a binary trait. We have relatively few models that explicitly model associations of genomic substitution rates with continuous phenotypes (Kowalczyk et al., 2019, 2020, 2022). Such continuous phenotypes likely better characterize many traits, and may provide additional power to link genotype and phenotype via phylogenetic trees.

## Data Availability

PhyloAcc and PhyloAcc-GT are open source software under the GNU General Public License (v3.0) and are freely available at https://phyloacc.github.io/. All input and output files for the analysis of the simulated data, ratite data, and mammal data as well as the scripts used to generate the figures in this manuscript are also available at https://github.com/phyloacc/Yan-etal-2022, with the exception of nucleotide alignments. These are available in the original PhyloAcc paper (Hu et al., 2019).

## Acknowledgements

This work was supported by NIH R01 HG011485-01. HY and JSL were also supported in part by NSF DMS-2015411. We would like to thank Taehee Lee, Patrick Gemmell, and Subir Shakya for discussion, and Nathan Weeks for computational advice. The computations in this paper were run on the FASRC Cannon cluster supported by the FAS Division of Science Research Computing Group at Harvard University.

## Supplemental Materials

### A Species trees for ratites and mammals

**Figure A.1:**
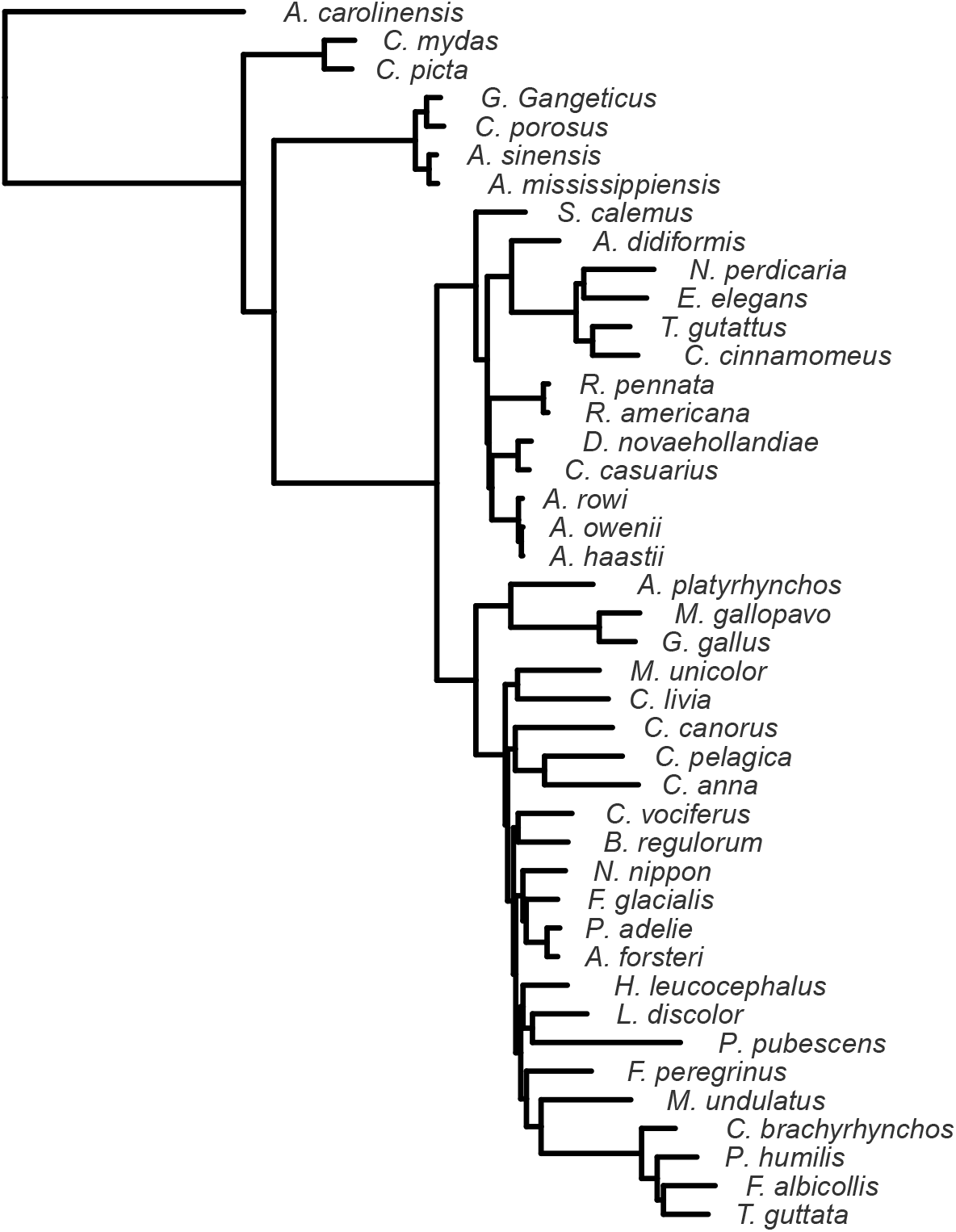
The full avian phylogeny used in this study, from Sackton et al. (2019) and Hu et al. (2019).

**Figure A.2:**
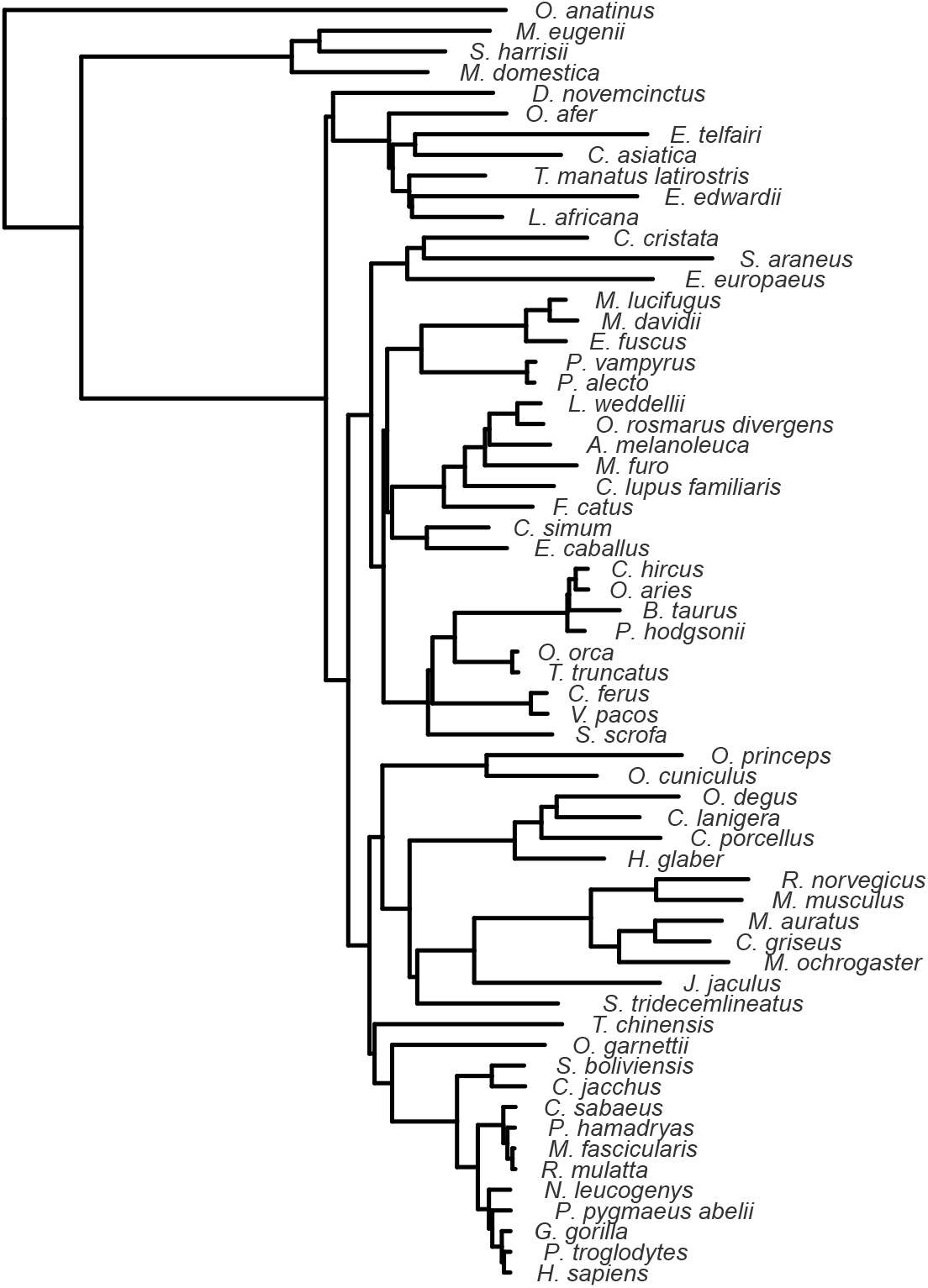
The full mammal phylogeny used in this study, from the UCSC 100-way vertebrate alignment Blanchette et al. (2004) and Hu et al. (2019).

### B Correctness of the proposed MCMC algorithm for gene tree inference

#### Case 1: Inferring branch length on a 2-leaf tree

To check the performance of the proposed MCMC algorithm for estimation of gene tree branch lengths, we first examine the simplest case: a 2-leaf tree. In this case, the two lineages only coalesce in the root species. There is no node above the root node on the gene tree to constrain our sampling. We set a hard threshold: 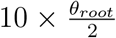 as the maximum height of the gene tree. Inferring the coalescent time is the same as inferring the position of the gene tree root. The Metropolis algorithm uses the uniform distribution centered at the current root node position as the proposal distribution. The step size is set as 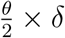, where *δ* ∈ [0.1, 5] is adaptive to ensure a reasonable acceptance rate. When the acceptance rate is too high, we will scale *δ* by a factor of 2; if the acceptance rate is too low, we will scale down *δ* to 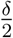. The proposal distribution is also constrained by the species tree and the upper limit of gene tree height that we set.

In this simple case, we can estimate some statistics of the posterior distribution, e.g., the posterior mean of the branch length, *l*, using numerical integration.

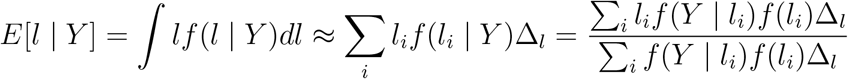

**Figure B.1:**
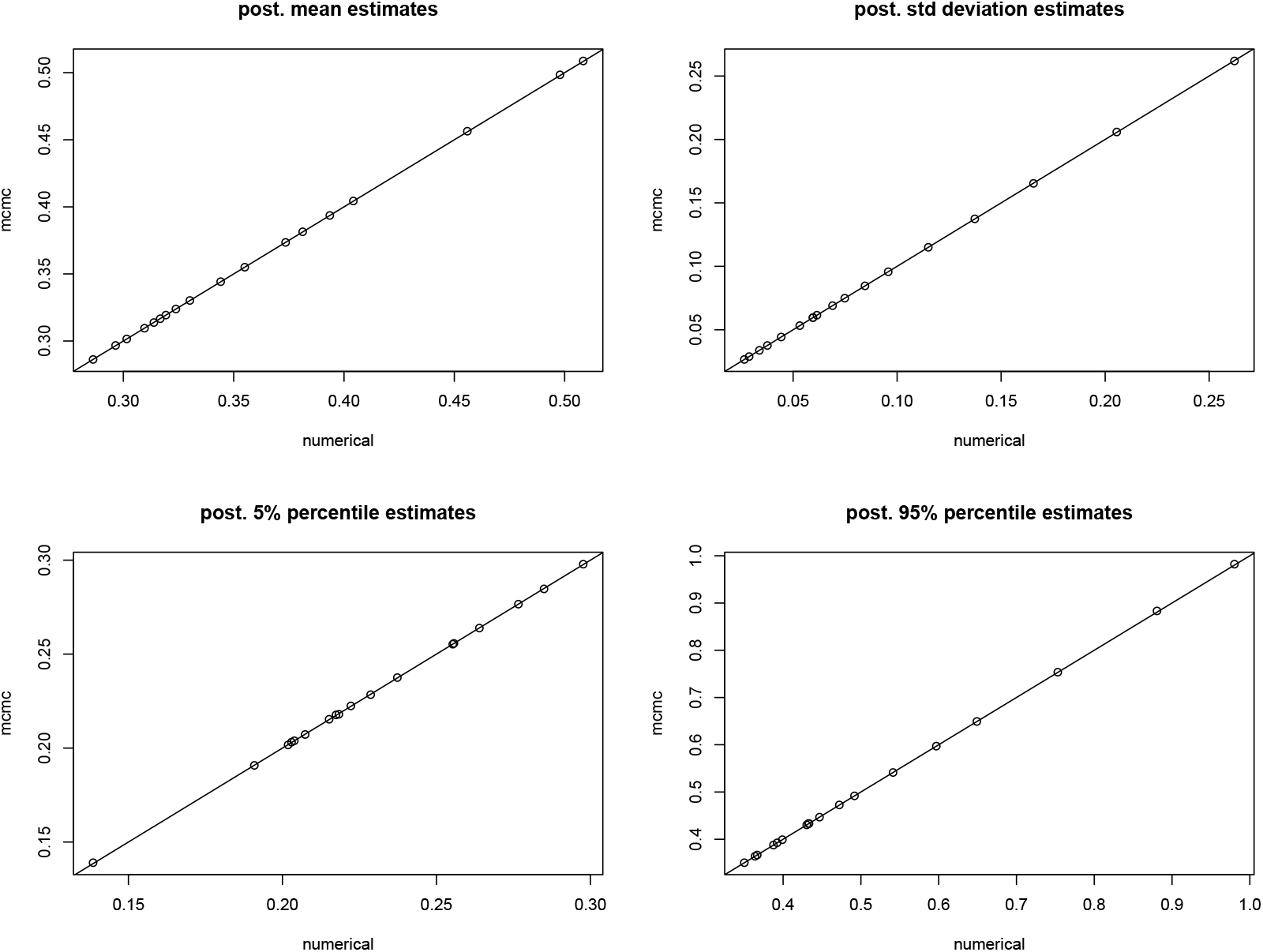
Comparison between our MCMC algorithm and numerical integration on some summary statistics of posterior distributions of coalescent time in the 2-leaf tree case. We ran the experiment under different number of base pairs ranging from 50 to 5000. We estimated posterior mean, standard deviation, tail probabilities: i.e. 5% quantile and 95% quantile using both MCMC sampling output and numerical integration. The x-axis represents results using numerical integration and the y-axis corresponds to the MCMC output. The line in each plot is *y* = *x*. For all four statistics, estimation results using the two methods fall almost perfectly along the *y* = *x* line.

**Figure B.2:**
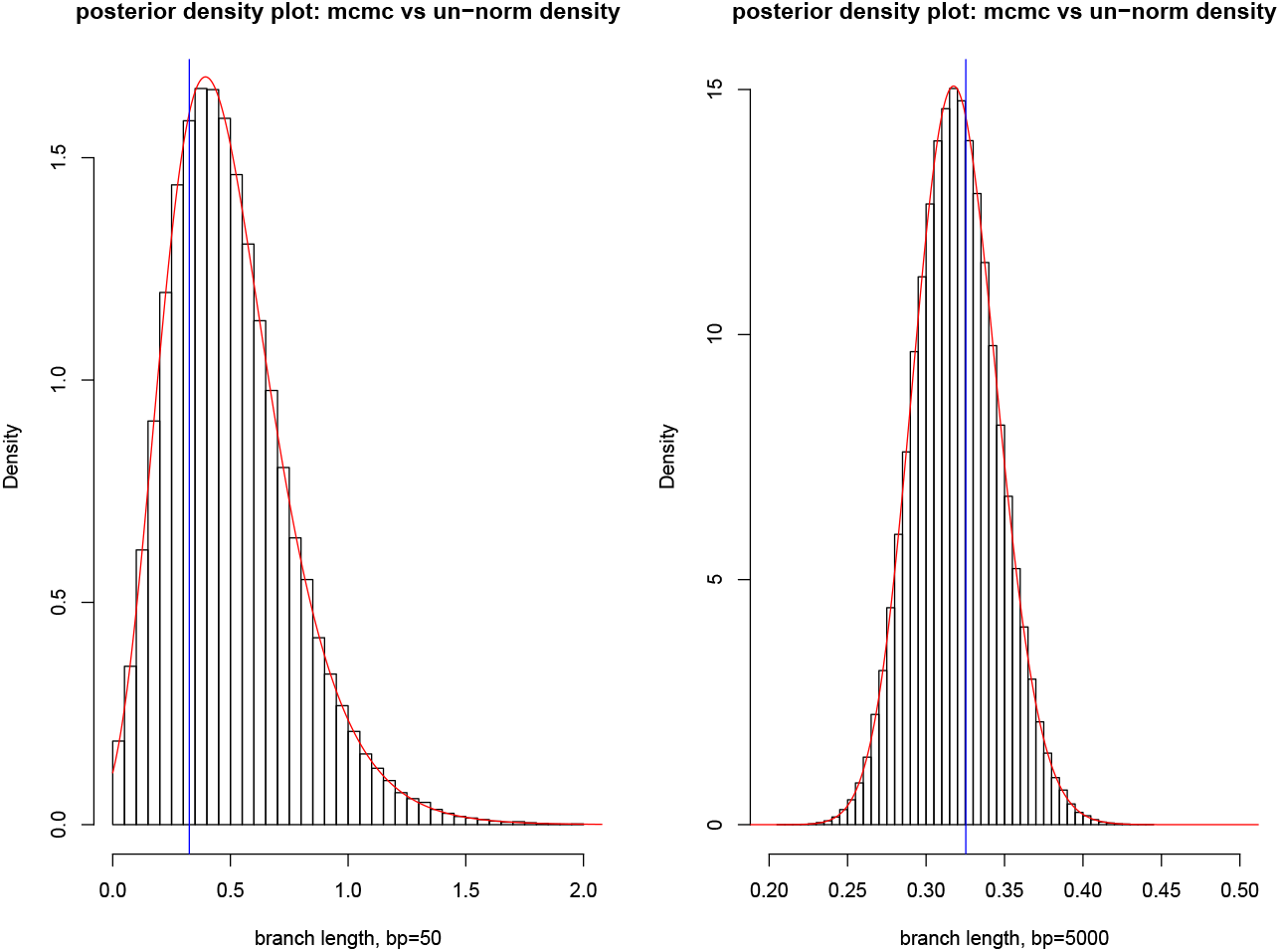
Posterior density plots in the 2-leaf tree case. The left plot corresponds to the posterior distribution of coalescent time with 50 base pairs, and the right plot is for 5000 base pairs. Histograms are based on MCMC sampling output. Red curves are plotted based on un-normalized posterior densities on grid points. Blue vertical lines are the true coalescent time. The red curves and histograms align very well, indicating our MCMC algorithm is sampling from the targeted posterior distributions. As the number of base pairs increases from 50 to 5000, the posterior distribution becomes more concentrated around the true coalescent time.

#### Case 2 - Inferring gene tree topology and branch lengths on a 3-leaf tree

To estimate the posterior probability of a gene tree topology, using Bayes’ theorem, we can write the posterior probability as:

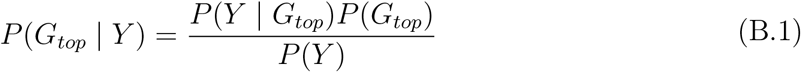

We estimate *P*(***Y***) by 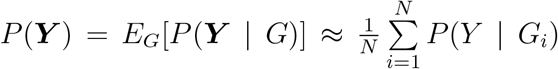, where a large number *N* of gene trees are simulated from its prior distribution. 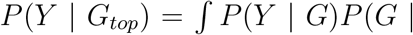 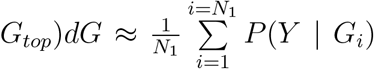, where *G_i_*, *i* = 1, · · ·, *N*_1_ are *N*_1_ prior trees with the sample topology *G_top_*. *P*(*G_top_*) can be estimated by the sampling proportion of the *N* prior trees with the particular topology denoted by *G_top_*. For a 3-leaf tree, the prior probability of each gene tree topology can be analytically calculated by integrating out all branch lengths. Let *G*_1_ denote the gene tree topology that is the same as the species tree (*T*) topology, and let *G*_2_ and *G*_3_ be the remaining two gene tree topologies.

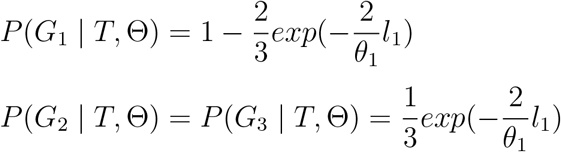

where *l*_1_ is the branch length on the species tree from the root to the first speciation event, and *θ*_1_ is the population size parameter of the species before the first speciation event. So Equation B.1 can be approximated by:

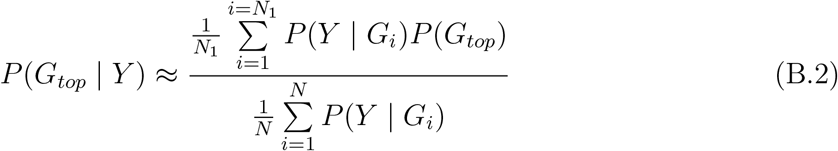

**Figure B.3:**
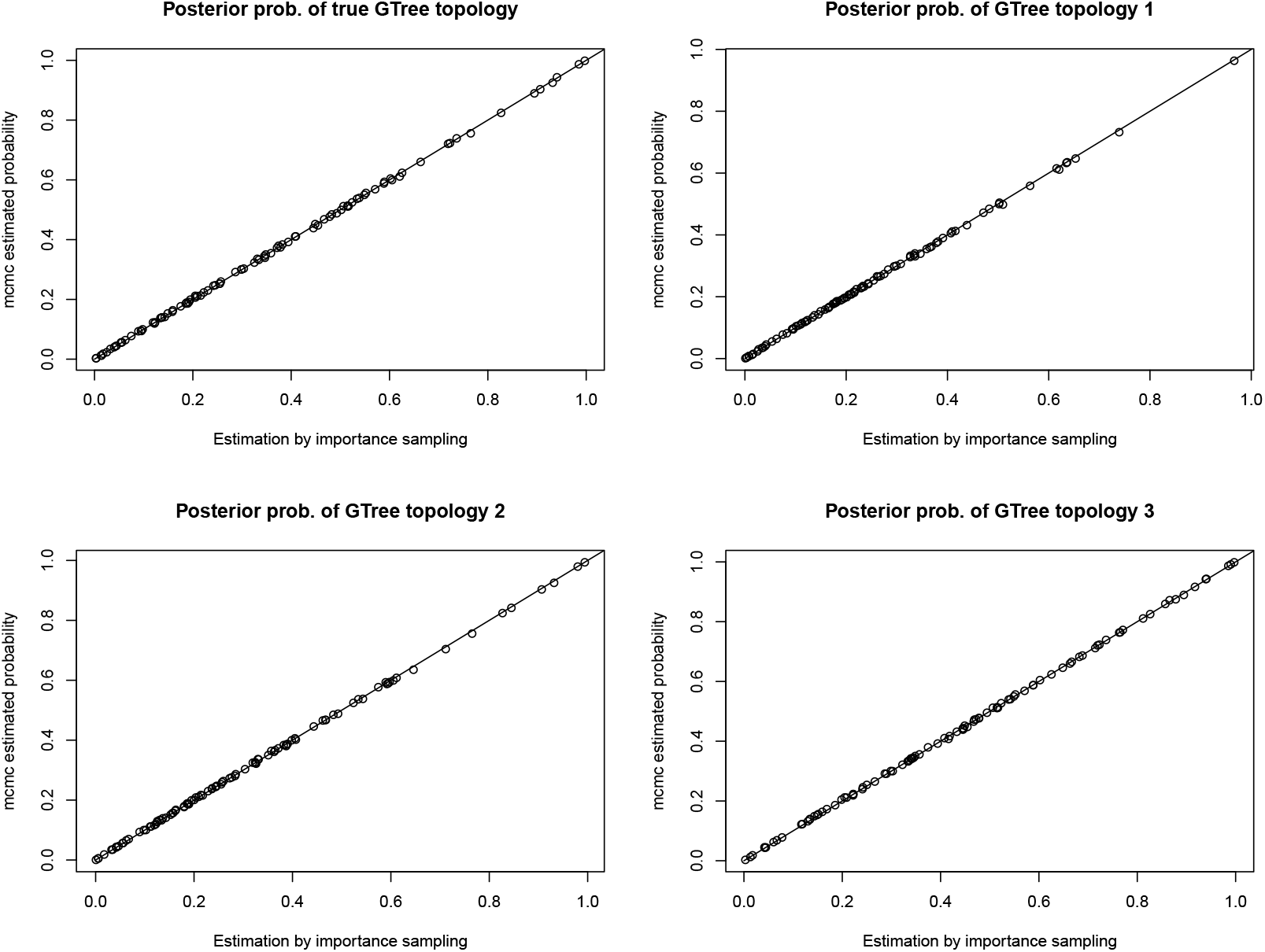
Estimating posterior probabilities of different gene tree topologies in the 3-leaf tree case. The x-axis corresponds to estimation using Equation B.2. The y-axis corresponds to the estimation using sampling frequencies of our MCMC algorithm. We generated 100 elements: i.e. 100 gene trees and base pairs based on each gene tree. The dots represent the estimation results for the 100 elements. The top-left plot compares estimation results for the posterior probability of the true gene tree topology. The remaining three plots are estimation results for the posterior probabilities of each of the three possible gene tree topologies. In all cases, the estimations by MCMC sampling frequency and by Equation B.2 are well aligned along the *y* = *x* line, indicating the correctness of our algorithm.

We also checked the correctness of our mcmc algorithm in inferring coalescent time under the 3-leaf tree case. The results are summarized in Figure B.4 and B.5.

**Figure B.4:**
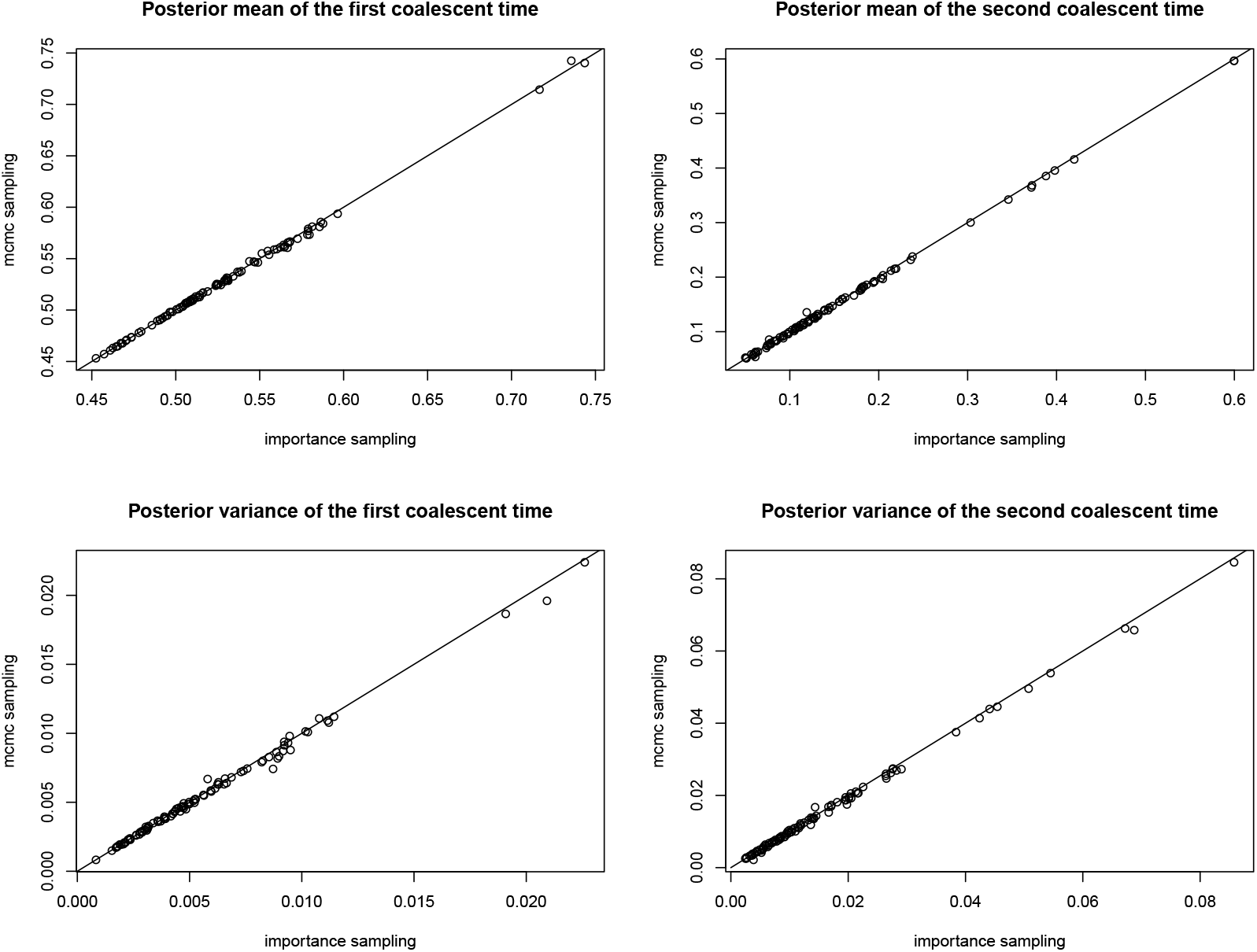
Correctness of the MCMC algorithm in inferring the posterior mean and variance of the two coalescent times in the 3-leaf tree case. Left plots correspond to the posterior mean and variance of the first coalescent time, and the plots on the right correspond to the second coalescent time. The x-axis corresponds to approximating the estimate by sampling branch lengths from the conditional prior distributions and approximating expectations by sample averages, similar to Equation (B.2). The two estimation methods are very close to each other.

**Figure B.5:**
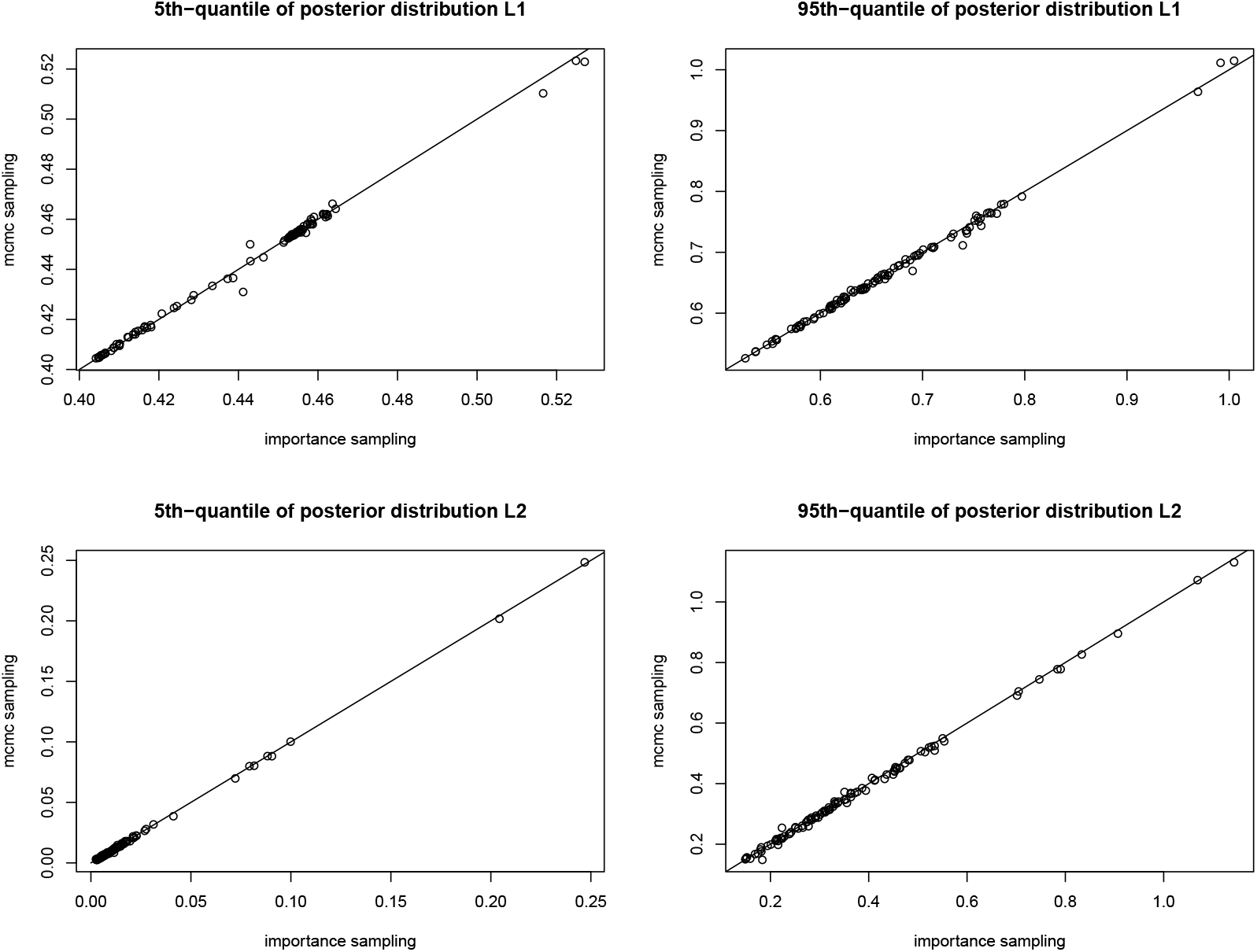
Correctness of the MCMC algorithm in estimating tail probabilities, i.e. 5% and 95% quantiles of the posterior distributions of the two coalescent times in the 3-leaf tree case. Upper plots are for the first coalescent time, and lower plots are for the second coalescent time. The two estimation methods gives very similar results, indicating the correctness of our MCMC algorithm.

### C Correctness of PhyloAcc-GT’s sampling algorithm for gene tree prior distribution

In PhyloAcc-GT we have implemented an algorithm to sample gene trees from their prior distribution conditioning on a species tree according to the multispeices coalescent model (Rannala and Yang, 2003). We use a simulation study to show that our sampling algorithm is correct. We show that several characteristics of the sampled gene trees match those of gene trees sampled from Phybase (Liu and Yu, 2010) in R given the sample species tree. We fix the species tree as:“((*A* : 0.05#0.01, *B* : 0.05#0.01) : 0.05#0.08, ((*C* : 0.06#0.01, *D* : 0.06#0.01) : 0.02#0.06, *E* : 0.08#0.01) : 0.02#0.05)#0.02;”, with topology plotted in Figure C.1.

**Figure C.1:**
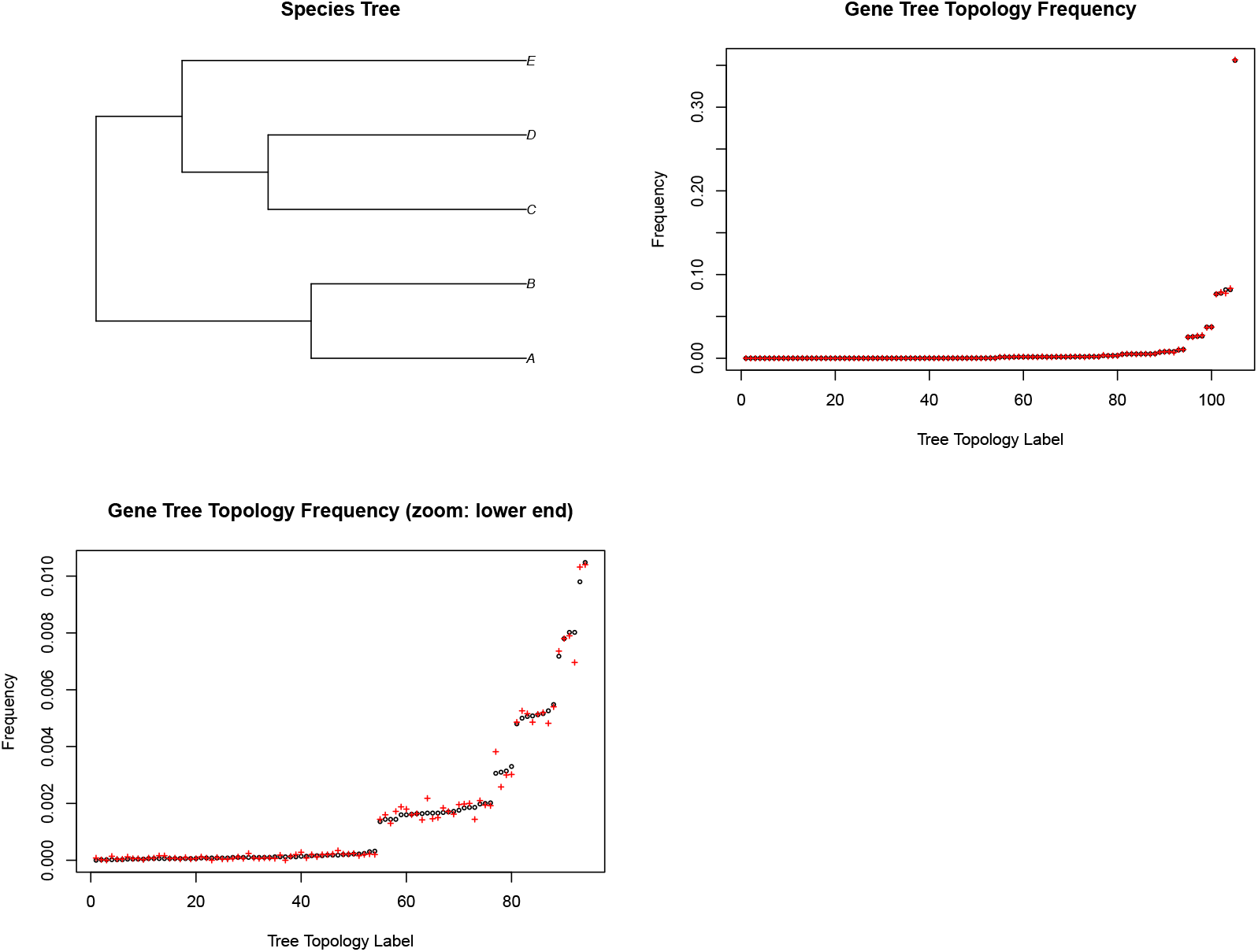
The species tree topology for simulation study in Section C, and gene tree sampling frequencies using PhyloAcc-GT and Phybase. Total number of samples is 50,000. The top right figure shows that sampling frequencies at various tree topology using PhyloAcc-GT (black dots) and Phybase (red plues) are very close to each other. The bottom left figure focuses on the low frequency end of the top right figure. With five extant species, the total number of rooted gene tree topologies is 105. The most frequently sampled gene tree topology matches the species tree topology in both algorithms. The corresponding frequencies are around 35%.

For each leaf node, we plot the histogram of its branch length. See Figure C.2.

**Figure C.2:**
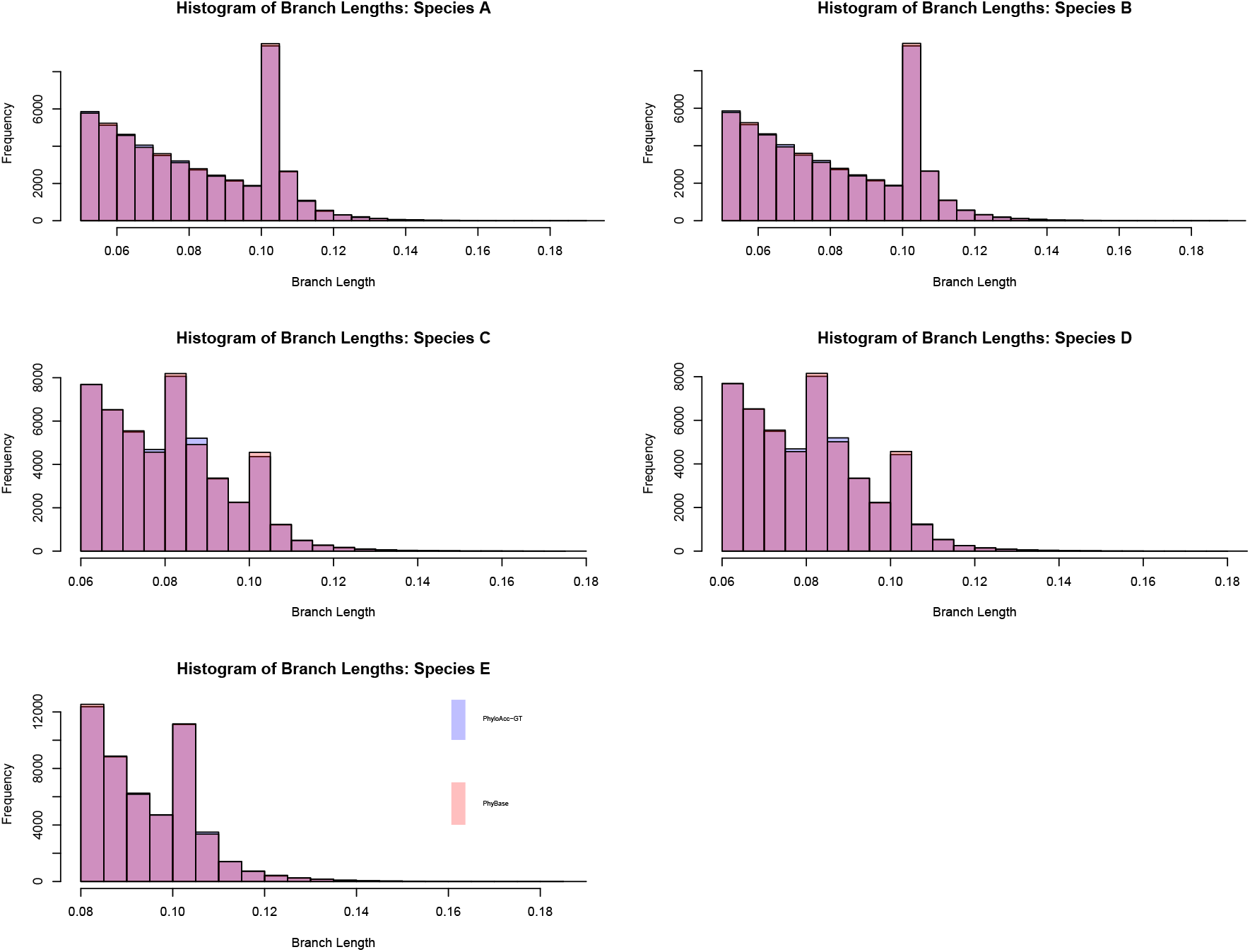
Overlapping histograms of sampled branch lengths of leaf nodes under PhyloAcc-GT and Phybase. For all 5 extant species, the sampling distributions of their branch lengths are very similar between PhyloAcc-GT and Phybase. The sampling distributions of the branch length of species A and the branch length of species B are very similar. The sampling distributions of the branch length of species C and the branch length of species D are also very similar. This result arises because in our samples, the genes in species A and B are most likely to coalesce first before coalescing with other lineages. A similar situation occurs with species C and D.

We also plot the histograms of the branch lengths of internal nodes of the most sampled gene tree topology in Figure C.3.

**Figure C.3:**
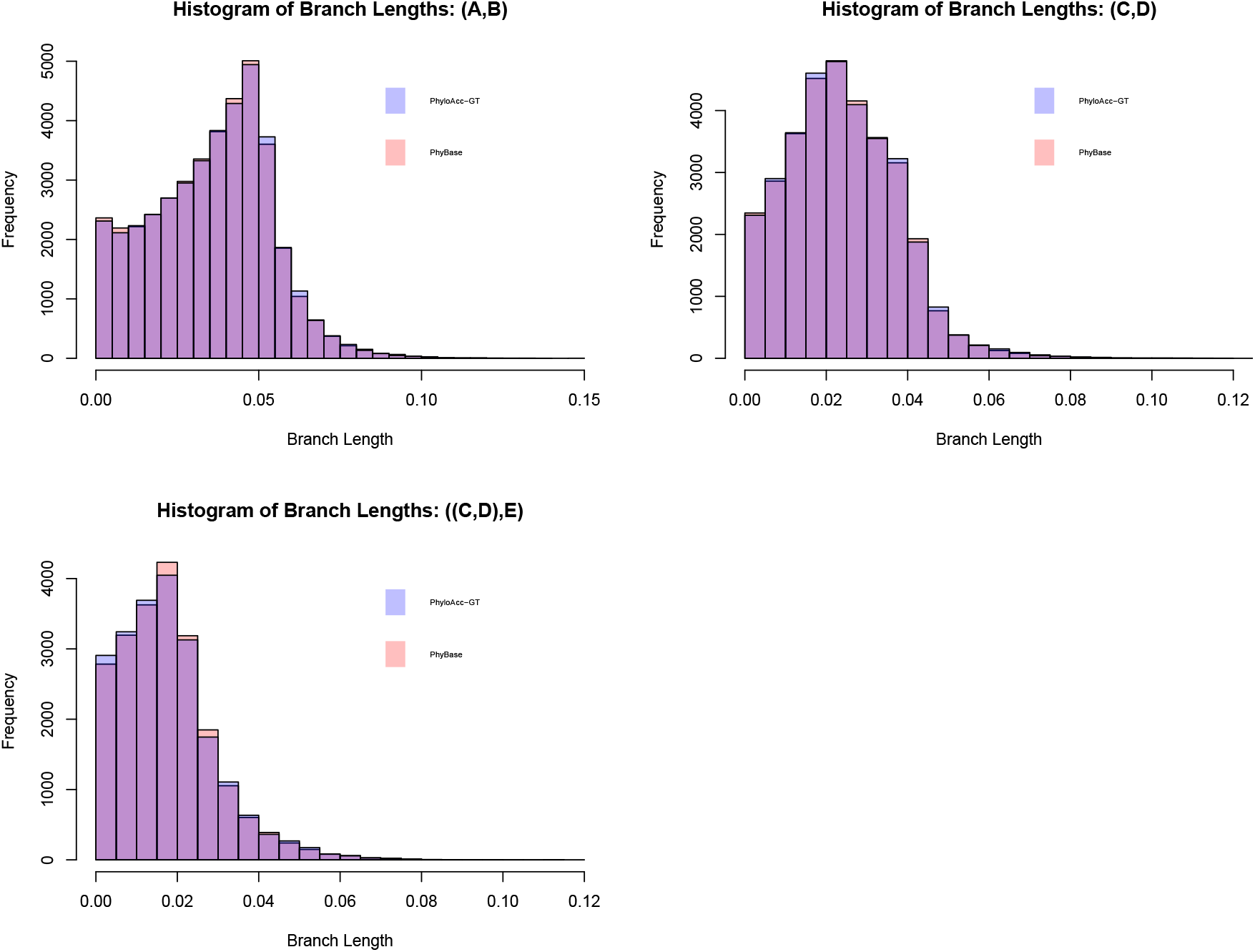
Overlapping histograms of some of the most frequently sampled internal nodes. The sampling distributions are similar between PhyloAcc-GT and Phybase.

Lastly we plot the histograms of some most frequently sampled internal nodes in Figure C.4.

**Figure C.4:**
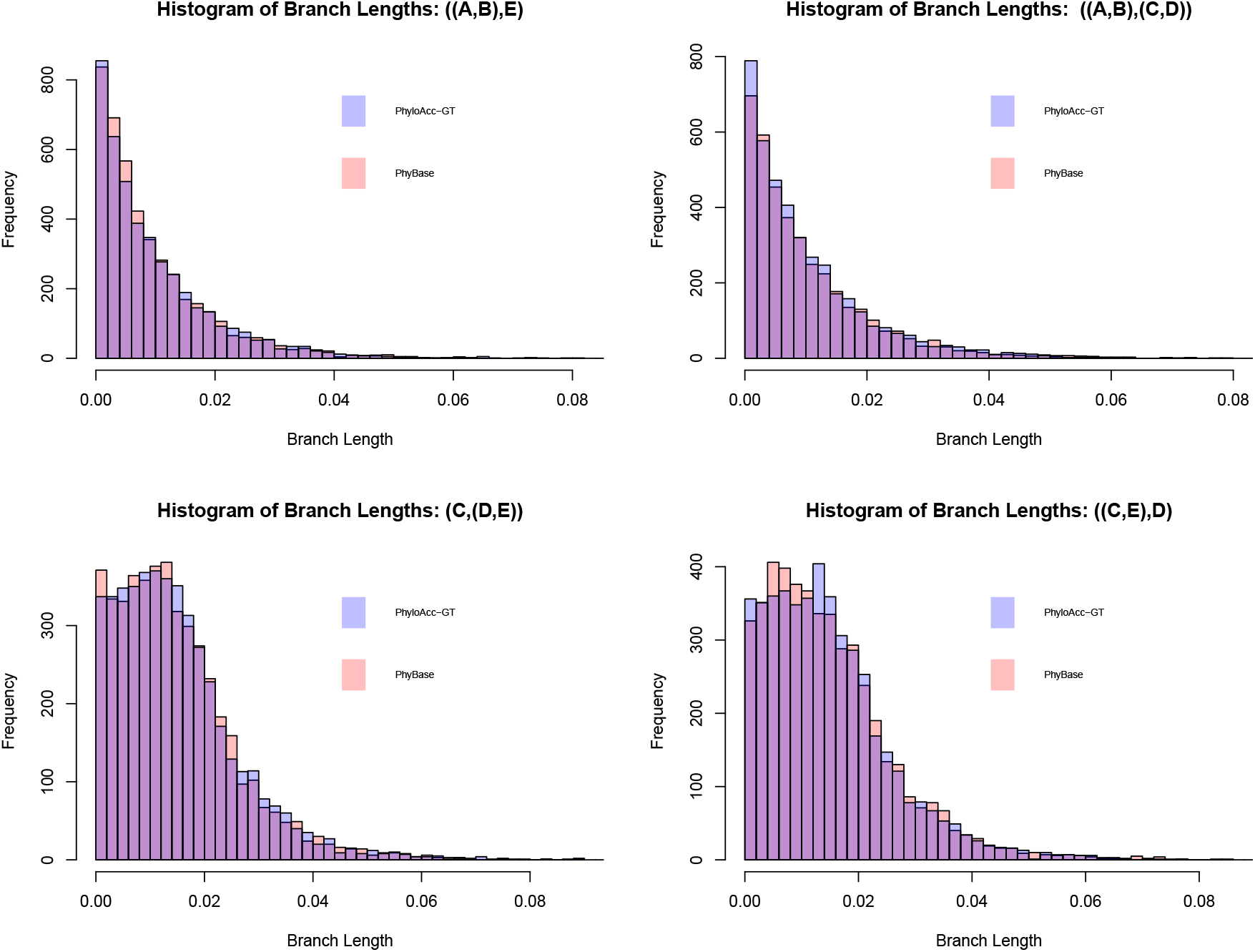
Overlapping histograms of some of the most frequently sampled internal nodes. The sampling distributions are similar between PhyloAcc-GT and Phybase.

### D Details of the Bayesian model

The prior state transition probability matrix Φ for ***Z*** is defined as: 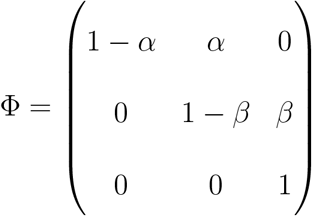. *α* represents the prior probability of an element becoming conserved from the background state in a lineage, and *β* is the prior probability of losing conservation. We put gamma priors on substitution rates and uniform priors on the hyperparameters *α* and *β*.

The prior distribution of a gene tree given the species tree is defined according to the multispecies coalescent model Rannala and Yang (2003), which we briefly review here. Let **Θ** = {*θ*_1_, · · ·, *θ_N_*} be population size parameters. For one species, *θ* = 4*N_s_μ*, where *μ* is the mutation rate per site per generation and *N_s_* is the population size. For each species, we record the coalescence events backwards in time until speciation. Suppose for an ancestral species *s* with branch length *t_s_*, there are *m_s_* sequences entering *s* at time 0, and *n_s_* leaving at time *t_s_*, with *n_s_* < *m_s_*. Let 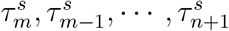 be the coalescent times for the time ordered (*m* − *n*)^*th*^ coalescent events, and 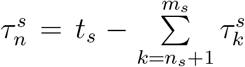 be the remaining time from the last coalescent event to the next speciation event. The prior density of a gene tree ***G*** is:

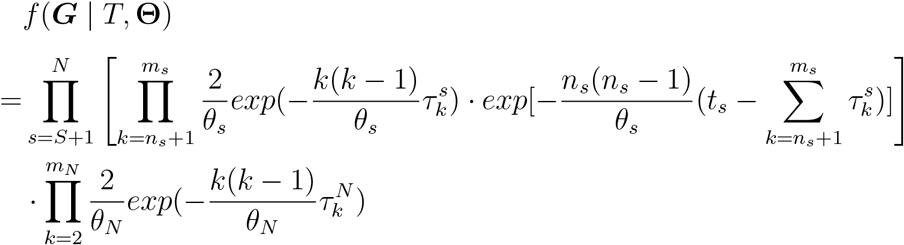

Given all parameters and latent variables, the complete data likelihood function is:

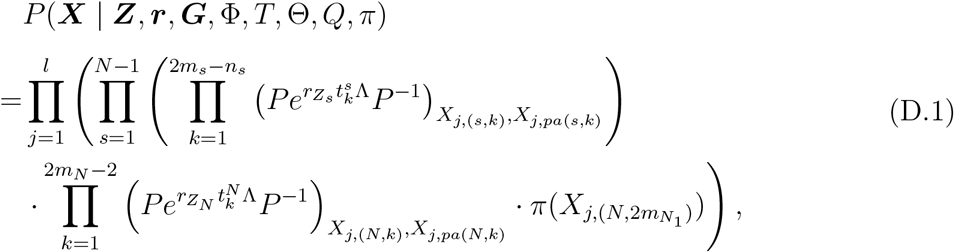

where ***X***_*j*,(*s*,·)_ contains base pair information at position *j* of the element for all sequences recorded in species *s*, and *X*_*j*,(*s,k*)_ for sequence *k* in *s*. 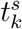 is the branch length from gene node (*s, k*) to *pa*((*s, k*)). *X*_*j*,(*s,k*)_ is the *j^th^* base pair in the *k^th^* sequence entering species *s* when *k* = 1, · · ·, *m_s_*, and is the *j^th^* base pair in gene node (*s, k*) generated by the (*k* − *m_s_*)^*th*^ coalescent event in species *s* when *k* = *m_s_* + 1, · · ·, 2*m_s_* − *n_s_*. The unnormalized posterior distribution is obtained by combining prior distributions with the full likelihood function.

### E Analyzing estimated rates

For PhyloAcc-GT and PhyloAcc, we also compare their estimated conserved rate and non-conserved rate under different patterns of acceleration. The result is shown in Figure E.5. For all cases, rates estimated by PhyloAcc-GT have higher correlations with the underlying true rates than rates estimated by PhyloAcc. PhyloAcc tends to overestimate rates, especially for the non-conserved rates, as can be seen in Figure E.6, E.7 and E.8. The overestimation is caused by ignoring gene tree heterogeneity due to incomplete lineage sorting, as well as the stationary distributions of nucleotide frequencies.

**Figure E.5:**
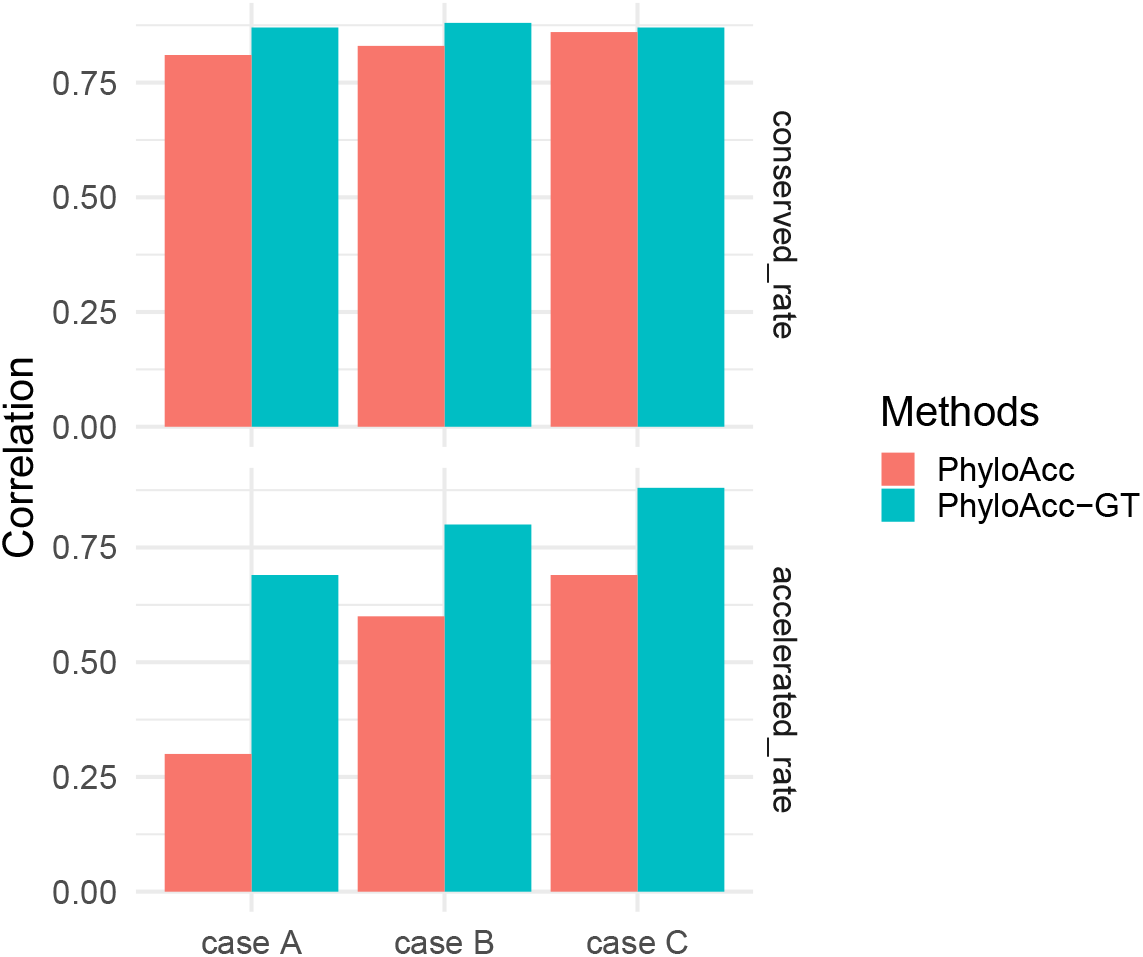
Comparing correlations between true rates and estimated ones by PhyloAcc-GT and PhyloAcc.

**Figure E.6:**
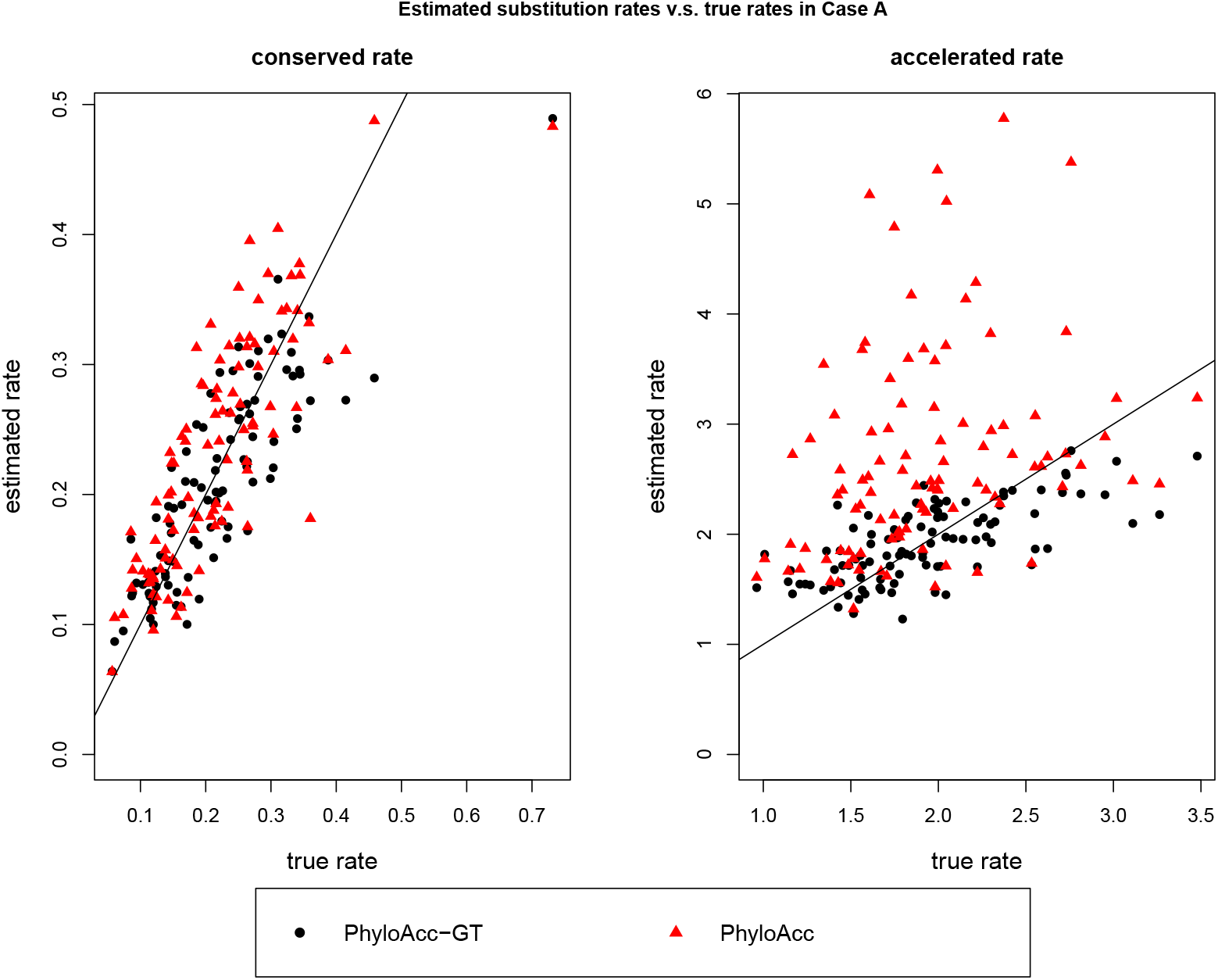
Comparing estimation of conserved and non-conserved rates by PhyloAcc-GT and PhyloAcc with sequences simulated with a single acceleration (2B). The line is Y=X.

**Figure E.7:**
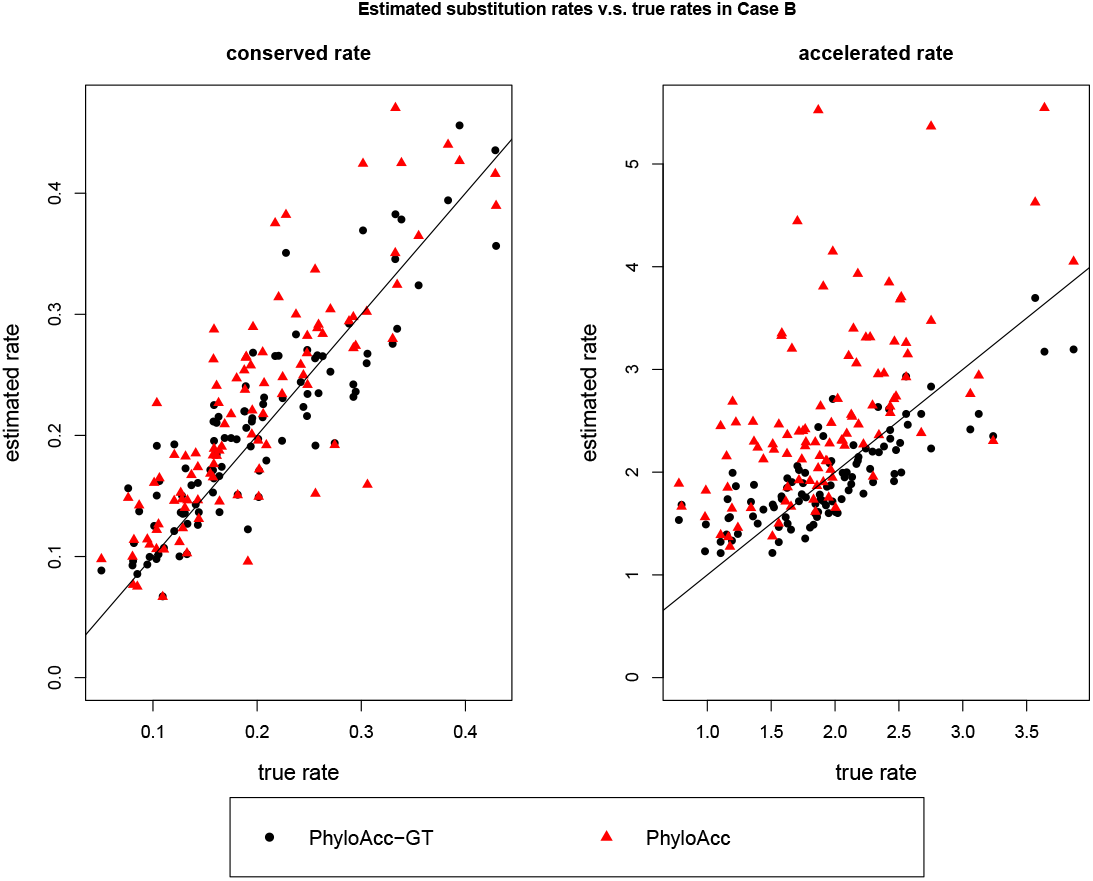
Comparing estimation of conserved and non-conserved rates by PhyloAcc-GT and PhyloAcc with sequences simulated with two independent (2C). The line is Y=X.

**Figure E.8:**
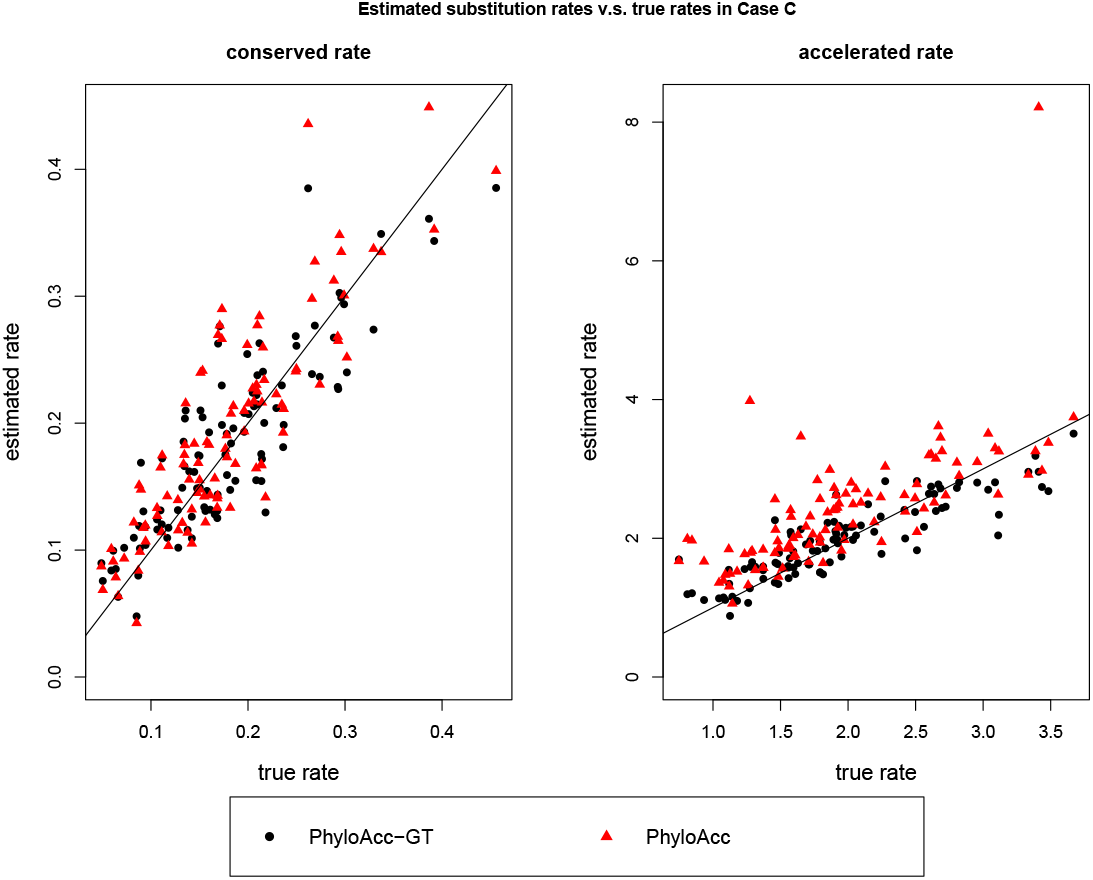
Comparing estimation of conserved and non-conserved rates by PhyloAcc-GT and PhyloAcc with sequences simulated with three independent (2D). The line is Y=X.

Our algorithm also gives good estimates of the frequencies of different nucleotides in the stationary distribution. We use the posterior mode as its point estimate. Plots of the estimated versus the true frequency of adenine in the stationary distribution are shown in Figure E.9 for two independent accelerations (2C). Relationships are very similar for the cases of a single acceleration 2B and three independent accelerations (2D), hence they are omitted. The correlations between the two are 0.927, 0.9 and 0.935 in the three simulation cases 2. Regressing the estimated *π_A_* against the true *π_A_* without an intercept term, the regression coefficient for two independent accelerations (2C) is 1.001, and 0.989 and 0.986 for a single acceleration 2B and three independent accelerations (2D respectively.

**Figure E.9:**
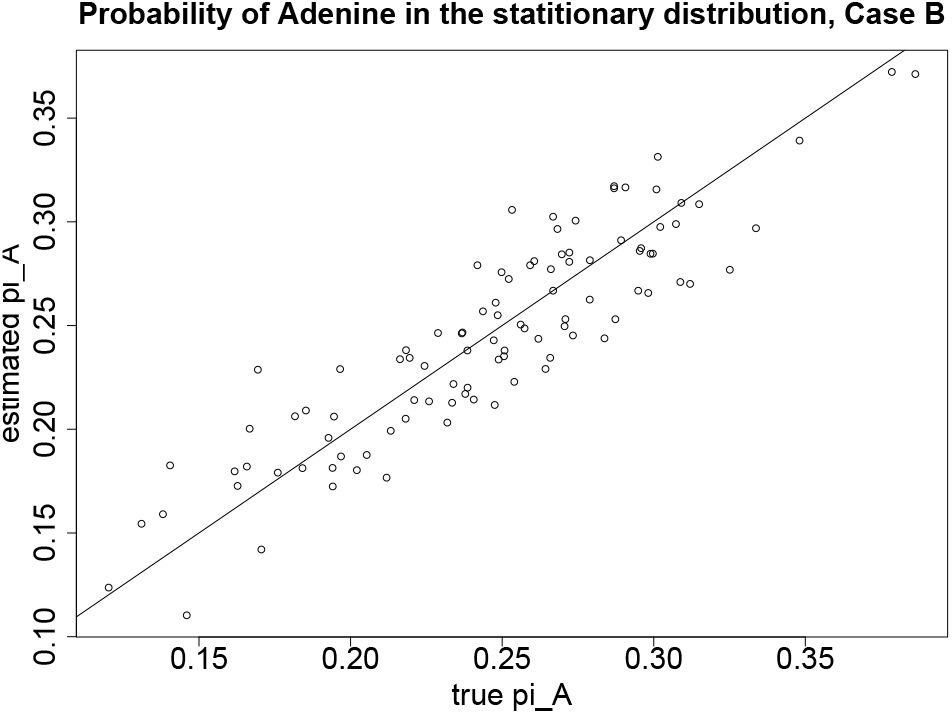
Estimated stationary probability of adenine v.s., the true probability with sequences simulated with a single acceleration (2B).

### F Modeling the substitution stationary distribution improves the estimation of substitution rates

The DNA nucleotide stationary distribution ***π*** directly affects the transition rate matrix *Q*. If *Q* is not correctly specified, it affects the estimation of substitution rates and potentially conservation states. In this section, we investigate the effect of ***π*** on model performance.

We use the same phylogeny as in Figure 2 A, and simulate DNA sequence from the null model, i.e., no branch is accelerated. We simulate 100 elements, having 200 base pairs each. Each element has its own ***π***. For 50 elememnts, we simulated 2*π_A_* ~ *Beta*(5, 5), and for the rest 50 elements, we simulated 2*π_A_* ~ *Beta*(10, 10). We have 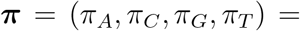 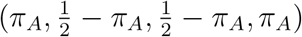. Conserved rates are simulated from *gamma*(5, 0.04). We run our algorithm for each element at 3 treatments of ***π***:

1. Treatment 1: fixing ***π*** at truth;
2. Treatment 2: fixing ***π*** at the value estimated from neutral sites, denoted by 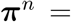 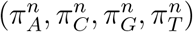;
3. Treatment 3: modeling the variation in ***π*** according to the Bayesian model in Section Methods in the main text.

When we designate the target group, for half of the elements we use sequences simulated with two independent accelerations (2C) and for the other half, we use sequences simulated with a single acceleration (2B).

Model selection accuracy is recorded in Table F.1. All three treatments are highly accurate in detecting no acceleration patterns along the phylogeny. Next, we check the posterior distributions of the conserved rate. We use the posterior distribution of *r*_1_ estimated estimated under Treatment 1 as a reference distribution. Figures F.1 and F.2 show that the posterior distributions are very close to the reference distribution whether we model ***π*** or not. Modeling ***π*** introduces slightly more variations in the upper tail probability of *r*_1_ as shown in the bottom left plot in Figure F.2.

**Table F.1:** Accuracy in model selection and *r_c_* estimation under the three models. Model selection accuracy is the percentage of cases the null model *M*_0_ is selected based on Bayes Factor cutoff at 1.

| Distribution to Simulate $\pi$ | Model | Model Selection Accuracy |
| --- | --- | --- |
| $Beta(10, 10)$ | 1. Fix $\pi$ at truth | 100% |
| | 2. Use $\pi^n$ | 100% |
| | 3. Estimate $\pi$ from data | 100% |
| $Beta(5, 5)$ | 1. Fix $\pi$ at truth | 96% |
| | 2. Use $\pi^n$ | 98% |
| | 3. Estimate $\pi$ from data | 98% |

**Figure F.1:**
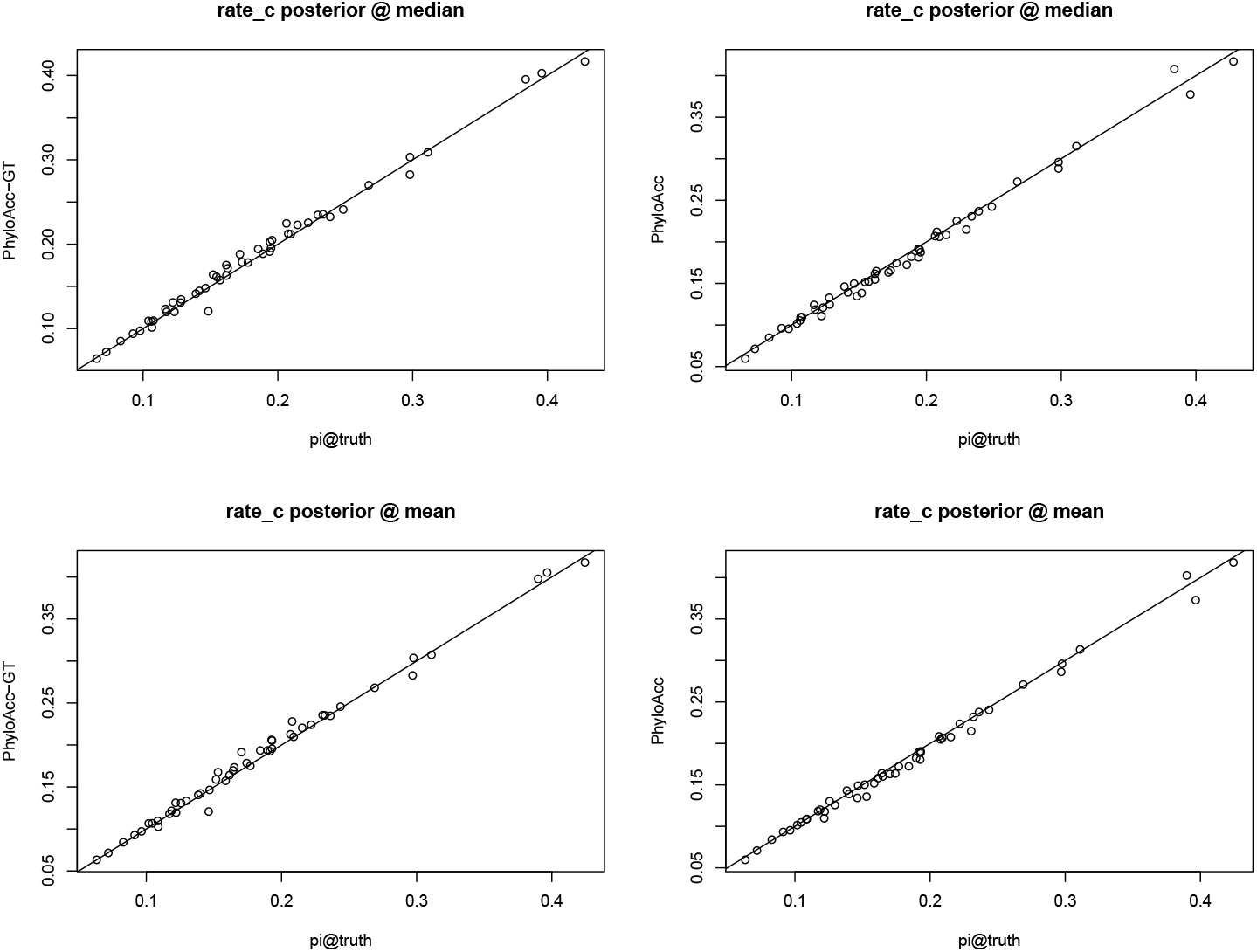
scatter plots comparing point estimates of *r_c_* using the three models.

**Figure F.2:**
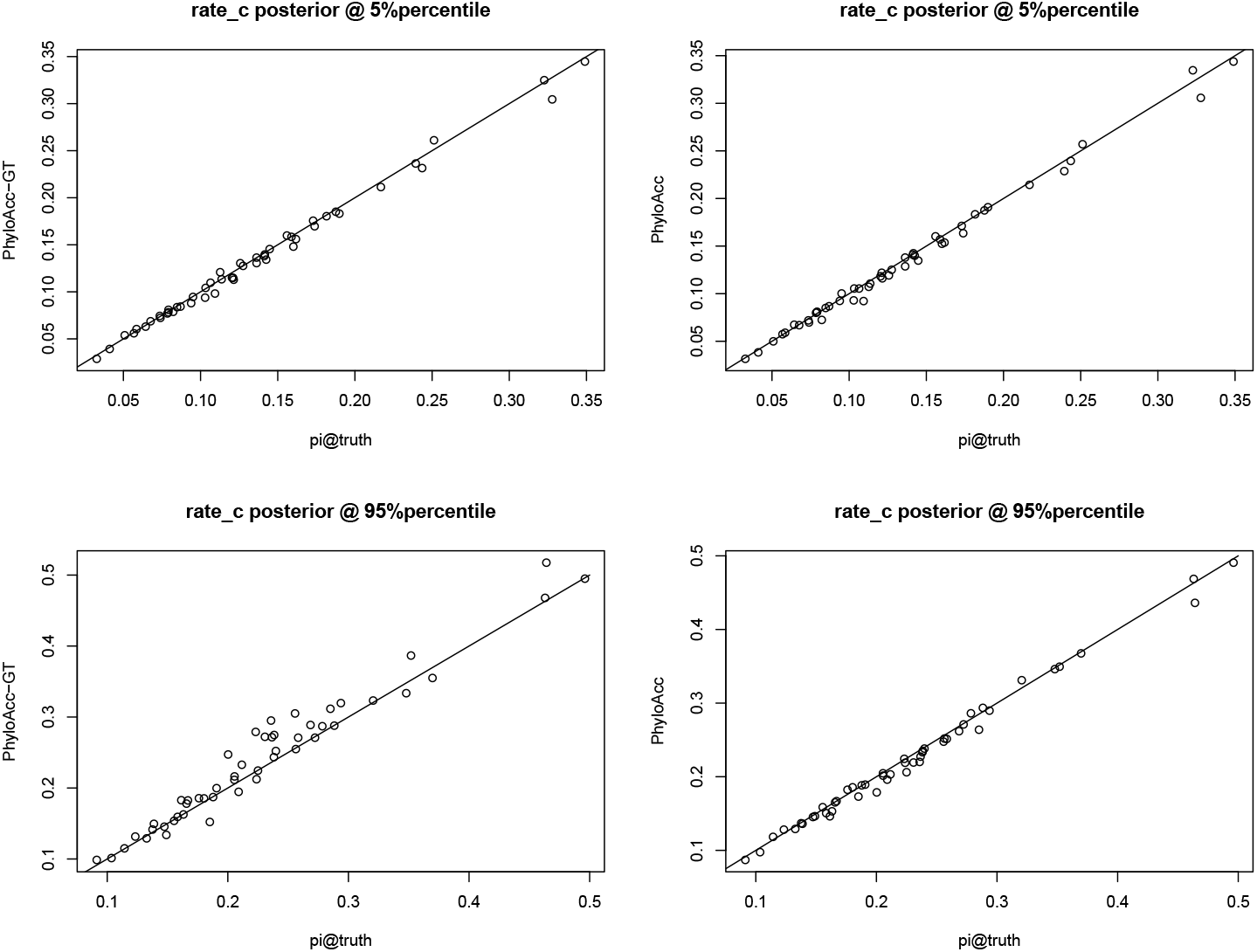
Scatter plots comparing tail behavior of the posterior distribution of *r_c_* using the three models.

The two Beta distributions we use to simulate *π_A_* + *π_T_* are centered at 0.5. Hence, the ***π***’s generated are likely to have balanced weights on all four nucleotides, and do not differ too much from ***π***^*n*^. The mean absolute difference between the simulated *π_A_*’s and 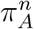 is 0.066. Since the differences are small, the rate matrix *Q* computed from ***π***^*n*^ also does not differ much from the true *Q*’s used to simulate the data. Results from the above study suggest that if ***π*** is misspecified by a small amount, it will not have an impact on inferring the posterior distributions of substitution rate and conservation state.

Next, we investigate whether modeling ***π*** will improve model performance from PhyloAcc when the input ***π*** value is far from the truth. We simulate 100 2*π_A_*s from two distributions: *Beta*(3, 1) and *Beta*(1, 4). *Beta*(3, 1) tends to produce ***π***s that put high weights on adenine and thymine, while *Beta*(1, 4) the opposite. We filter out values of *π_A_* that are either greater than 0.4 or less than 0.1, which gives us 56 cases. Using these unbalanced ***π***’s we generate data and test our models. When applying PhyloAcc, we input a ***π*** that put most weight on cytosine and guanine if the true ***π*** is highly concentrated on adenine and thymine, or the other way around.

In this extreme case, PhyloAcc still achieves 100% model selection accuracy, and the accuracy is 94.4% for PhyloAcc-GT. Both are accurate in identifying 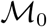 as the correct model. However, PhyloAcc underestimates the substitution rate as shown in Figure F.3, while PhyloAcc-GT can still accurately inference the substitution rate. If the data set is generated with a large *π_A_* value, most of the base-pair positions will show A or T across extant species. However, since the input *π_A_* is very small when running PhyloAcc, the sequences are more likely to transit from A and T to C and G. To observe the high frequency of A and T and high similarities among sequences, PhyloAcc has to infer that DNA substitution will be highly conserved.

**Figure F.3:**
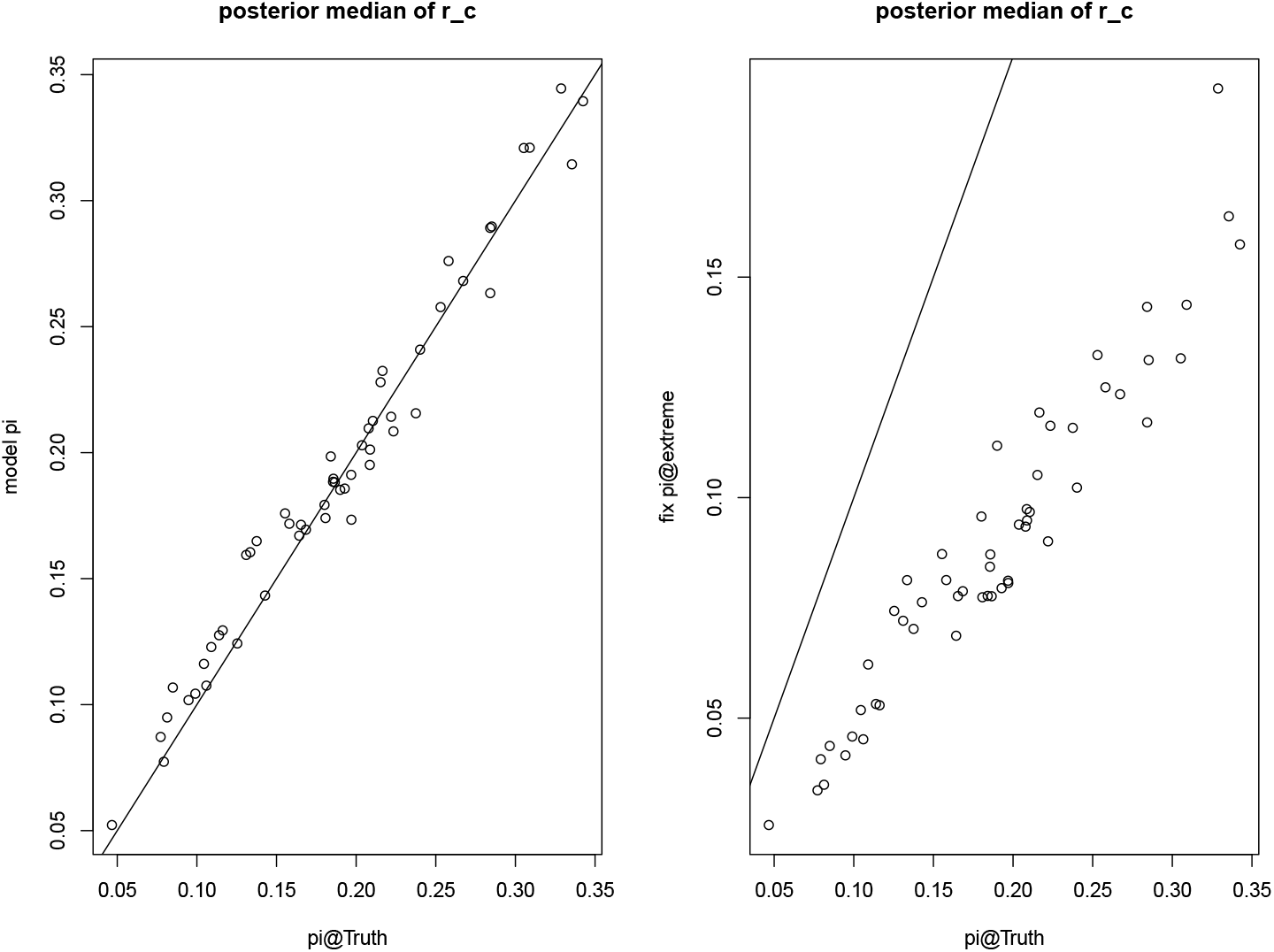
Comparing posterior mediums of the substitution rate from the three models

### G Some Analyses of Posterior Gene trees and Patterns of Acceleration for Elements of Interest in the Avian Dataset

In this section, we analyze posterior gene trees for element mCE1358939 and mCE1419808 that favor model 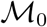 under PhyloAcc-GT. For mCE1358939, with PhyloAcc, Southern cassowary, Little spotted kiwi and Great spotted kiwi are estimated to be in the accelerated state with posterior probability of acceleration being 0.75, 0.85 and 0.56 respectively. It is likely that the acceleration in the two kiwis occurred in their parent species (*P* (*Z* = 2 | ***Y***) = 0.52). Under PhyloAcc-GT, the four species are still the top four species that are likely to have experienced rate accelerations under 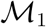, but only Southern cassowary and Little spotted kiwi have posterior probabilities of acceleration exceeding 0.5. Under 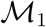, the gene tree at the posterior mode places the Rhea clade directly under Ostrich, and (Southern Cassowary, Emu) becomes the sibling branch of (Moa, Tinamous). The same tree topology is also the most likely topology under model 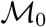. However, there are increases in the estimated gene tree branch lengths for the four branches under 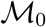, and most non-accelerated branches are shorter under 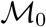 than under 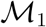.

**Figure G.1:**
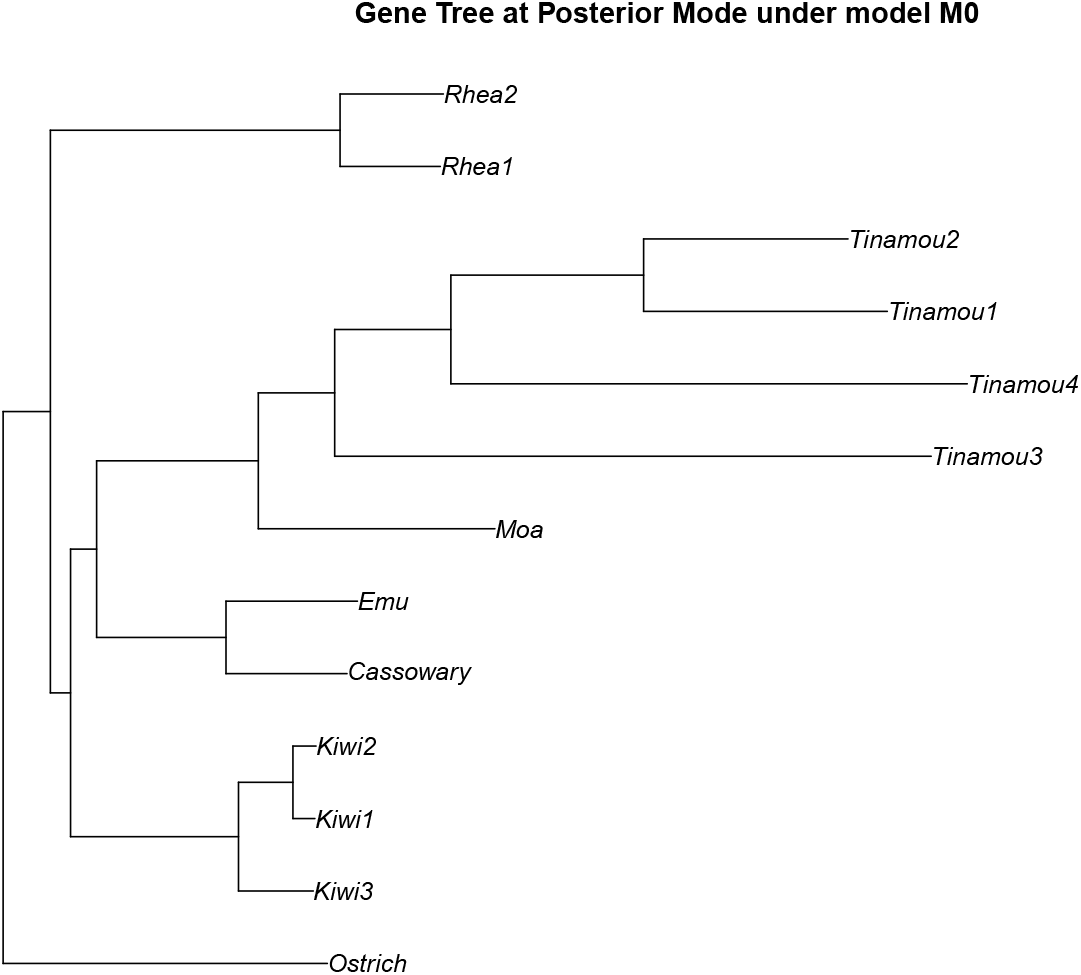
The gene tree topology at posterior mode under model 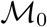 for element mCE1358939.

Using PhyloAcc, mCE1419808 is estimated to have experienced strong rate accelerations in Ostrich (*P* (*Z* = 2 | ***Y***) = 1), followed by Great spotted kiwi and Little spotted kiwi (*P* (*Z* = 2 | ***Y***) = 0.56 for both). Using PhyloAcc-GT, the gene tree topology among ratites at posterior mode under 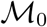 and 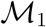 are both the same as the species tree topology. However, the gene tree branch lengths differ from those of the species tree, resulting in different patterns of acceleration. In the posterior mode of the gene tree under model 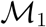, the estimated branch length for Ostrich is 8% longer than the corresponding length on the species tree. As a result, the estimated posterior probability of acceleration in Ostrich reduces to 0.5 under PhyloAcc-GT. On the other hand, posterior probabilities of acceleration are greater in (Great spotted kiwi, Little spotted kiwi), (Greater rhea, Lesser rhea), and (Cassowary, Emu) under PhyloAcc-GT compared to the estimated probabilities using PhyloAcc, because gene tree branch lengths are estimated to be shorter than species tree branch lengths. Although some branches are estimated to have rate accelerated under 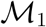, and the ratite tree topology at the posterior mode are the same under 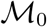 and 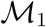, after marginalizing over the gene tree, the data supports model 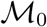 the most.

### H Additional Figures from Simulation Studies

**Figure H.1:**
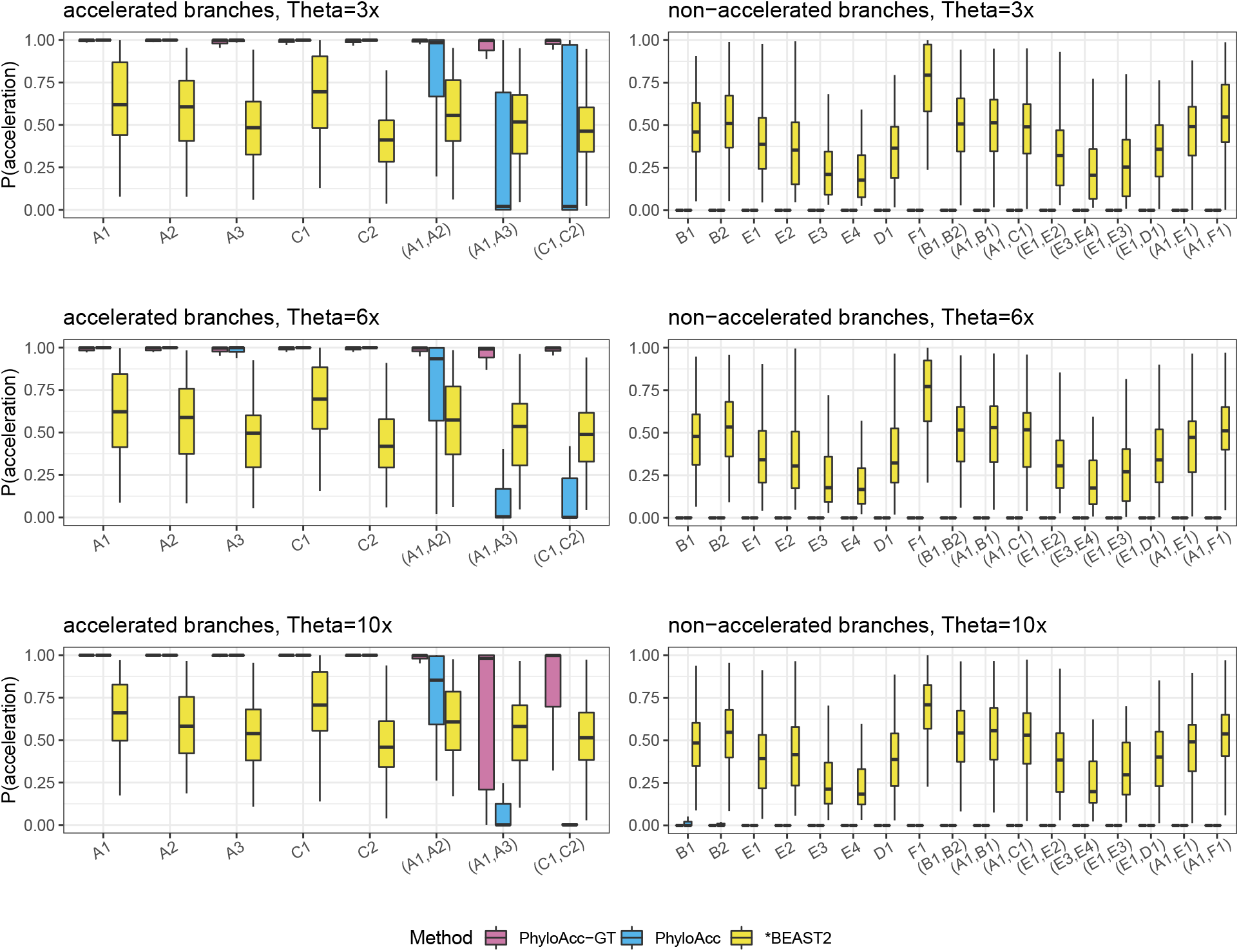
comparing *P* (*Z* = 2 | ***Y***) using PhyloAcc-GT, PhyloAcc and *BEAST2 under the two independent accelerations case (2C) as **Θ** increases. Left plots correspond to truly accelerated branches, whereas plots on the right correspond to non-accelerated branches. We multiply all *θ* values by 3, 6 or 10, shown in top, middle and bottom rows respectively.

**Figure H.2:**
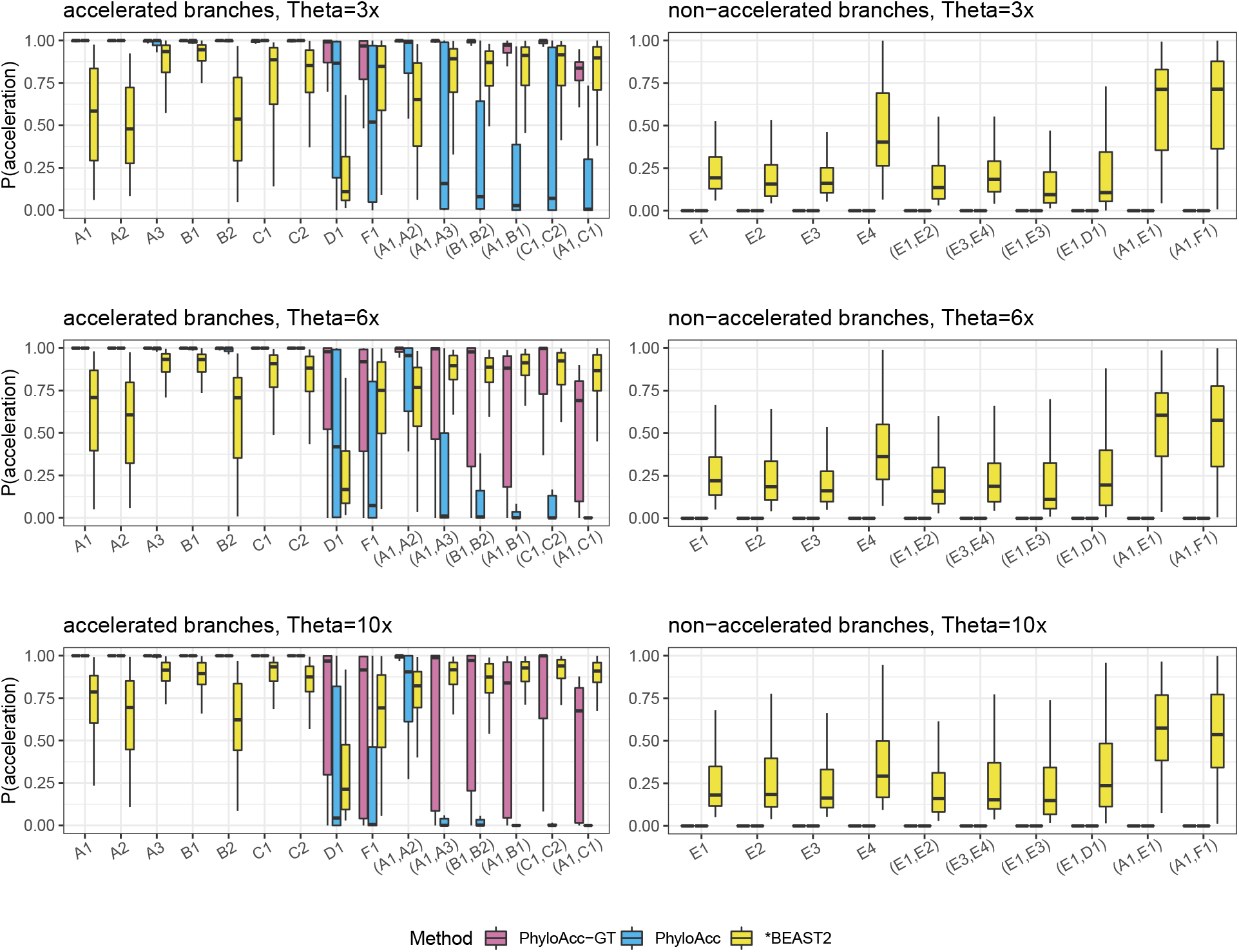
comparing *P* (*Z* = 2 | ***Y***) using PhyloAcc-GT, PhyloAcc and *BEAST2 under the three independent accelerations case (2D) as **Θ**increases. Left plots correspond to truly accelerated branches, whereas plots on the right correspond to non-accelerated branches. We multiply all *θ* values by 3, 6 or 10, shown in top, middle and bottom rows respectively.

